# Optimising the production of dsRNA biocontrols in microbial systems using multiple transcriptional terminators

**DOI:** 10.1101/2024.02.22.581520

**Authors:** Sebastian J. Ross, Gareth R. Owen, James Hough, Annelies Philips, Wendy Maddelein, John Ray, Peter M. Kilby, Mark J. Dickman

**Affiliations:** Department of Chemical and Biological Engineering, University of Sheffield, S1 3JD, UK; Syngenta Innovation Center, Gent, 9052, Belgium; Syngenta, Jealott’s Hill International Research Centre, Bracknell, Berkshire, RG42 6EY, UK

**Keywords:** dsRNA, RNAi, RNA biocontrols, *E.coli*, transcriptional terminators, mass spectrometry, agricultural biotechnology

## Abstract

Crop pests and pathogens annually cause over $100 billion in global crop damage, with insects consuming 5-20% of major grain crops. Current crop pest and disease control strategies rely on insecticidal and fungicidal sprays, plant genetic resistance, transgenes and agricultural practices. dsRNA is emerging as a novel sustainable method of plant protection as an alternative to traditional chemical pesticides. Successful commercialisation of dsRNA based biocontrols requires the economical production of large quantities of dsRNA combined with suitable delivery methods to ensure RNAi efficacy against the target pest. In this study, we have optimised the design of plasmid DNA constructs to produce dsRNA biocontrols in *E. coli*, by employing a wide range of alternative synthetic transcriptional terminators prior to measurement of dsRNA yield. We demonstrate that a 7.8-fold increase of dsRNA was achieved using triple synthetic transcriptional terminators within a dual T7 dsRNA production system compared to the absence of transcriptional terminators. Moreover, our data demonstrates that batch fermentation production dsRNA using multiple transcriptional terminators is scalable and generates significantly higher yields of dsRNA generated in the absence of transcriptional terminators at both small-scale batch culture and large-scale fermentation. In addition, we show that application of these dsRNA biocontrols expressed in *E. coli* cells results in increased insect mortality. Finally, novel mass spectrometry analysis was performed to determine the precise sites of transcriptional termination at the different transcriptional terminators providing important further mechanistic insight.

## Introduction

A rapidly growing global population presents a significant challenge to meet the increasing demand for food in the coming years. It has been estimated that by 2050, crop production may need to increase by 50% to satisfy demand (Mcloughlin et al., 2018). Approximately 40% of total crop production is lost, primarily through pests and pathogens (Wytinck et al., 2020), resulting in an annual loss of over $100 billion. This figure could escalate to $540 billion due to the spread of invasive pathogens and pests (Niehl et al., 2018). Biotic factors alone are responsible for 17-30% of global annual yield losses in the five major food crops (Wytinck et al., 2020). Insects, in particular, are responsible for consuming 5-20% of major grain crops, and it is projected that the damage caused by them will increase by 10-25% with every global temperature degree increment (Deutsch et al., 2018).

Current crop pest control strategies rely on insecticidal and fungicidal sprays and plant genetic resistance and/or traditional agricultural practices. There is a growing demand for innovative, sustainable approaches to crop protection driven by an increasing population, climate-driven pest range expansion, community and regulatory demands, and pest resistance to traditional agro-chemicals (Savary et al., 2012).

RNA interference (RNAi) is emerging as an important tool for the development of novel RNA-based sustainable insect management strategies (Dalaisón-Fuentes et al., 2022; Katoch et al., 2013). Application of double-stranded RNA (dsRNA) or endogenous dsRNA expression in genetically engineered plants has been utilised for the sequence specific degradation of targeted mRNA in crop pests (Baum & Roberts, 2014; Mao et al., 2011; Yan et al., 2020).

Non-transformative methods, as defined in the review by Hough et al., (2022), are used to produce exogenous dsRNA, these include *in vitro* transcription (IVT), microbial expression in bacteria or fungi, and cell-free synthesis, offering rapid and controlled production of dsRNA, suitable for experimental research and scalable quantities. In contrast, transformative methods involving genetically modified (GM) plants allow continuous dsRNA production, beneficial for applications like pest-resistant crops, reducing pesticide use and enhancing yields. However, GM crop methods raise concerns regarding gene flow, and regulatory complexities. Both methods have their advantages and limitations, with each of the production methods providing specific modes of action (Christiaens et al., 2020; Hough et al., 2022)

The production of dsRNA in *E. coli* has predominantly used the RNase III deficient strain HT115 (DE3) (Ahn et al., 2019; Bento et al., 2020; García et al., 2015; Hull & Timmons, 2004; Meng et al., 2020; Nwokeoji et al., 2016; Ongvarrasopone et al., 2008; Posiri et al., 2013). Initial plasmid DNA constructs consisted of two opposing convergent T7 promoters, flanking the dsRNA sequence. The L4440 plasmid, which lacks transcriptional terminators, first designed, and used by Timmons and Fire in 1998, has been widely employed for microbial expression of dsRNA and subsequent RNAi studies (Timmons et al., 2001). Adaptations to the L4440 construct using T7 terminators outside of the T7 promoters suggested that this diminished the effectiveness of initiating RNAi via feeding in *C. elegans* (Kamath et al., 2000). In contrast, Sturm et al., (2018) demonstrated that the efficiency of RNAi was improved by the incorporation of T7 terminators.

During RNAi studies in *A. thaliana*, dual terminators have been employed (James A Baum & James K Roberts, 2014). In a study by Chen et al., (2019), a construct was developed to produce hpRNA using double terminators at the end of a single T7 promoter sense-loop-antisense sequence. The hpRNA system was compared to a conventional T7 system flanked by dual terminators, and a twofold increase in dsRNA production was observed with the hpRNA system.

Previous work has focussed on the development of more efficient T7 RNA polymerase terminators that enable higher protein expression *in vivo* with associated lower metabolic burden or for insulation of expression modules with multigene expression plasmids (Du et al., 2009; Du et al., 2012; Mairhofer et al., 2015). In these studies, novel synthetic terminators were developed and included the use of class I terminators, tandem class II terminators and cassettes with class I and class II terminators. More recently novel chimeric and compact terminators incorporating Class II pause sequences within Class I-hairpins have also been developed that exhibit strong termination efficiency *in vivo* and *in vivo* (Calvopina-Chavez et al., 2022).

In this study we designed a range of novel plasmid constructs aimed at optimising the production of dsRNA in *E. coli* utilizing multiple sets of synthetic T7 RNA polymerase terminators. We investigated the effect of synthetic terminators on the total dsRNA yield, quality of *in vivo* produced dsRNA biocontrols and RNAi efficacy *in vivo* using insect mortality assays. Furthermore, we determined the termination efficiency of these synthetic terminators within convergent T7 RNA polymerase promoter constructs for dsRNA production and provide further mechanistic insight in the T7 RNA polymerase termination using novel mass spectrometry approaches.

## Materials and Methods

### Chemicals and reagents

Ampicillin sodium salt, tetracycline hydrochloride, isopropyl β-D-1-thiogalactopyranoside (IPTG) ≥99%, sodium dodecyl sulphate (SDS) and sodium chloride (NaCl) were sourced from Sigma Aldrich. LB Miller media was sourced from Sigma Aldrich. Agarose gels were prepared using either molecular grade agarose (Appleton) or UltraPure agarose (Invitrogen). Gels were run using 1X Tris-acetate EDTA (TAE) buffer (Sigma Aldrich). Samples were stained with either Midori green direct dye (Geneflow) or Ethidium bromide (Alfa Aesar). RNA samples were loaded with Novex™ TBE-Urea Sample Buffer (2X) (Thermo Fisher scientific). *In vitro* transcription (IVT) eactions were performed with HiScribe™ T7 High Yield RNA Synthesis Kit (New England Biolabs). Template DNA was degraded with TURBO DNase (Thermo Fisher Scientific). IVT-produced RNA samples were purified using a Monarch® RNA Cleanup Kit (New England Biolabs). ssRNA during RNA extractions was degraded with RNase T1 (Thermo Scientific). UltraPure™ Phenol:Chloroform:Isoamyl Alcohol (25:24:1, v/v) was sourced from Thermo Fisher Scientific. To aid DNA pellet visualisation GlycoBlue™ Coprecipitant (Thermo Fisher Scientific) was used PCR reactions were performed using KAPA2G Fast Hotstart Readymix (Merck) and purified using Monarch® PCR & DNA Cleanup Kit (New England Biolabs). Gel fragments were extracted using GeneJet Gel extraction kit (Thermo Fisher Scientific). Ligations were performed using Quick Ligation™ Kit (New England Biolabs). HPLC and LC-MS mobile phases were prepared using ≥99.0% (GC) dibutylamine (Sigma-Aldrich), 99.5+% 1,1,1,3,3,3-hexaflouro-2-propanol (Thermo Scientific), Triethylammonium acetate pH 7.4 (Sigma-Aldrich), UHPLC-MS grade acetonitrile (Thermo Scientific), and UHPLC-MS grade water (Thermo Scientific).

### Biological sources

The HT115(DE3) *E. coli* strain was acquired from Jealott’s Hill International Research Centre, Syngenta, UK, while the DH5α strain of *E. coli* was obtained from New England Biolabs. Synthetic genes and DNA fragments were procured from either GeneArt® Gene Synthesis (Thermo Fisher Scientific) or Genewiz (Azenta Life Sciences). Table 1 contains a list of all plasmids used in this study.

**Table 1.**
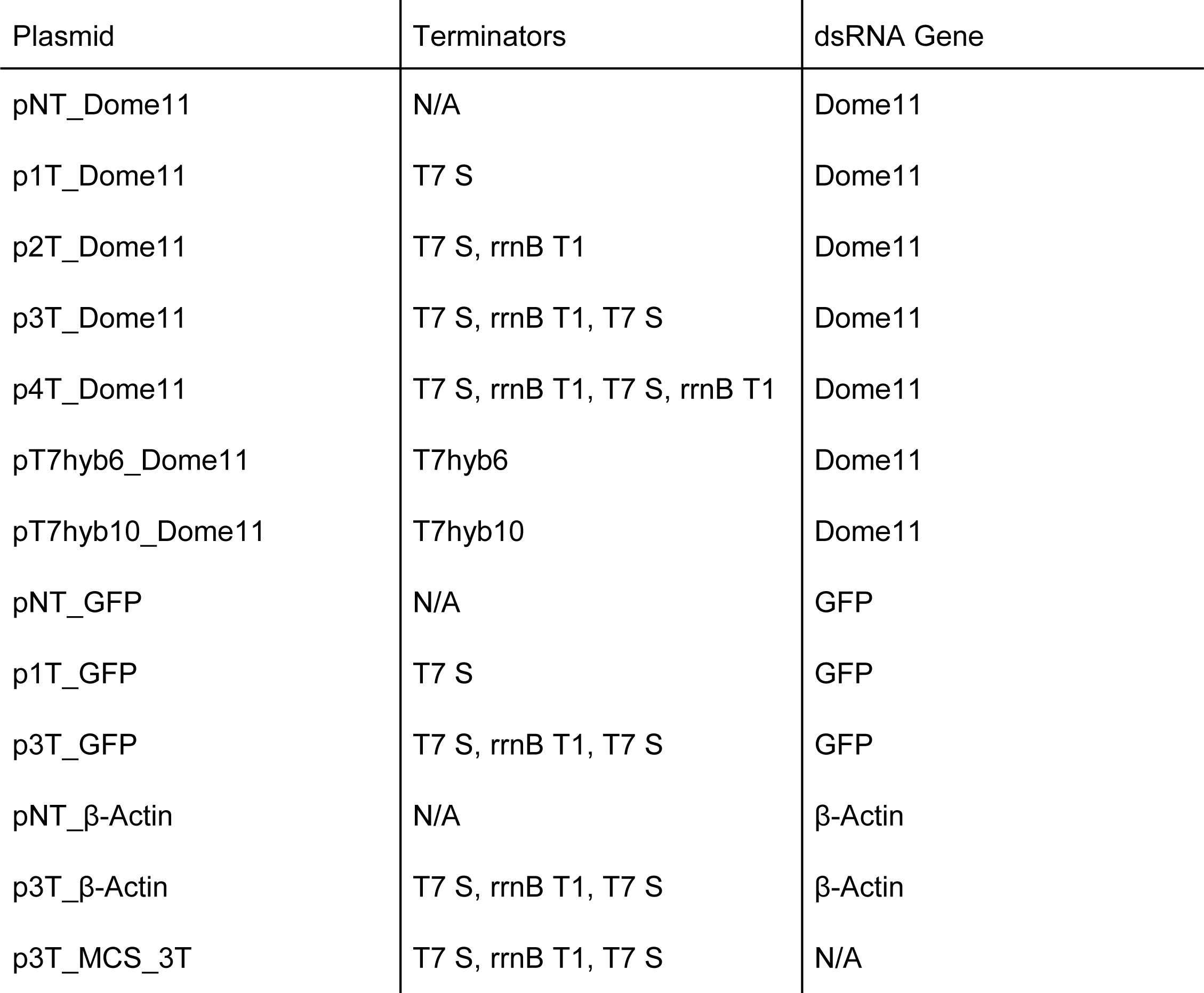

| Plasmid | Terminators | dsRNA Gene |
| --- | --- | --- |
| pNT_Dome11 | N/A | Dome11 |
| p1T_Dome11 | T7 S | Dome11 |
| p2T_Dome11 | T7 S, rrnB T1 | Dome11 |
| p3T_Dome11 | T7 S, rrnB T1, T7 S | Dome11 |
| p4T_Dome11 | T7 S, rrnB T1, T7 S, rrnB T1 | Dome11 |
| pT7hyb6_Dome11 | T7hyb6 | Dome11 |
| pT7hyb10_Dome11 | T7hyb10 | Dome11 |
| pNT_GFP | N/A | GFP |
| p1T_GFP | T7 S | GFP |
| p3T_GFP | T7 S, rrnB T1, T7 S | GFP |
| pNT_β-Actin | N/A | β-Actin |
| p3T_β-Actin | T7 S, rrnB T1, T7 S | β-Actin |
| p3T_MCS_3T | T7 S, rrnB T1, T7 S | N/A |

### *E. coli* cell growth and inductions

#### Small-scale shake flasks

A single transformed colony was inoculated into 5 ml of LB Miller media (Sigma) supplemented with 100 μg/mL ampicillin and 10 μg/mL tetracycline, then grown overnight at 37 °C, 250 rpm. After overnight incubation, 2 ml of the *E. coli* cells were fed into 50 mL of LB media containing the aforementioned antibiotic concentrations and incubated at 37°C, 250 rpm, until the OD_600_ nm reached ∼0.4-0.6. Induction was initiated by addition of IPTG to attain a final concentration of 0.1 mM, cells were then incubated for an additional 4 hours at 37 °C, 250 rpm.

#### Large-scale fermentation

Large-scale production of Dome11 dsRNA was conducted in a 1 L Multiflors 1 bioreactor (Infors HT). The starter culture was prepared via inoculation of 50 ml of LB media containing 100 μg/mL ampicillin, then grown overnight for 15 hours at 30 °C, 250 rpm. The starter culture was used to inoculate 50 ml of LB media, which was grown for 8 hours at 30 °C, 250 rpm. Fermentation inoculation was performed at a 1:20 (v/v) in 750 ml of LCM fermentation media containing 100 μg/mL ampicillin. Fermentation inoculate was grown at 24 °C overnight, and increased to 37 °C the following morning. Samples were induced upon reaching ∼10 OD_600_ nm with 1mM IPTG. Induction was performed at 37 °C for 3.5 hours, followed by an overnight growth at 20 °C. Dissolved oxygen (DO) was maintained at 30% of air saturation and pH was maintained at 6.8 by using 2M H_2_SO_4_ and 4M NaOH.

#### RNA extraction and purification

RNA purifications were performed as previously described by Nwokeoji et al., (2016) with minor modifications. Cells were aliquoted in quantities of 1 × 10^9^ cells, this was calculated using the online tool provided by Agilent (available at agilent.com/store/biocalculators/calcODBacterial.jsp), followed by incubation with preheated lysis buffer at 95 °C for 5 minutes, followed by the addition of 170 μL of 5 M NaCl and further incubation on ice for 5 minutes. The lysate was then centrifuged for 7 minutes, the supernatant collected and 200 μL of 100% isopropanol added before purification using a silica-membrane column (Geneflow). The wash steps followed the RNAswift protocol, and the RNA was eluted twice with preheated 50 μL of Ambion Nuclease-free water (Invitrogen) at 95 °C for 2 minutes each time. For certain investigations, a modified extraction to remove ssRNA was performed via the addition of 1 μl of 1/50 diluted RNase T1 (1000 U/µL) (Thermo Scientific) to the lysate before isopropanol addition and incubation for 45 minutes at 37 °C. RNA was analysed for both quantity and contamination using a Nanodrop 2000c UV spectrophotometer (Thermo Fisher Scientific). To quantify total dsRNA, an absorbance factor of 46.52 μg/mL per A_260_ was used as per Nwokeoji et al., (2017).

#### Agarose Gel electrophoresis

Molecular grade agarose (Appleton) or UltraPure agarose (Invitrogen) was used to prepare the agarose gels, which were then run using 1X Tris-acetate EDTA (TAE) buffer (40 mM Tris (pH 7.6), 20 mM acetic acid, and 1 mM ethylenediaminetetraacetic acid (EDTA)) with varying percentages based on the specific sample analysis required. E-Gel™ EX Agarose Gels (Thermo Scientific) were also utilized for sample analysis. To visualize the samples, either Midori green direct dye (Geneflow) or ethidium bromide (Alfa Aesar) staining was applied. DNA samples were mixed with Trilink loading buffer (Thermo Fisher Scientific), while RNA samples were mixed with Novex™ TBE-Urea Sample Buffer (2X) (Thermo Fisher Scientific). Samples loaded onto EX Agarose Gels were mixed with the E-Gel sample buffer (1X) (Thermo Scientific). Agarose gels were visualized under blue light using a Fas-DIGI illuminator (Geneflow). For sizing purposes, the following ladders were utilized: GeneRuler 1 kb plus (Thermo Scientific), dsRNA ladder (NEB), ssRNA ladder (NEB), RiboRuler Low Range RNA Ladder (Thermo Scientific), and RiboRuler High Range RNA Ladder (Thermo Scientific).

#### In vitro transcription

PCR templates or linearised plasmids were utilized for *in vitro* transcription. The linearised templates were prepared by restriction enzyme digestion followed by phenol/chloroform ethanol precipitation. The HiScribe™ T7 High Yield RNA Synthesis Kit (New England Biolabs) was used for in vitro transcription according to the manufacturer’s instructions, with samples incubated at 37 °C for 2 hours unless specified otherwise. Degradation of DNA templates was performed via the addition of 1 μL of TURBO DNase (2U/μL) (Thermo Fisher Scientific) at 37 °C for 20 minutes. RNA samples were purified using the Monarch® RNA Cleanup Kit (New England Biolabs) following the manufacturer’s instructions.

#### Phenol/chloroform ethanol precipitation

UltraPure™ Phenol:Chloroform:Isoamyl Alcohol (25:24:1, v/v) (Thermo Fisher Scientific) was added to samples at a volume ratio of 1:1, followed by brief vortexing and centrifugation at 15,200 rpm for 10-15 minutes at 4 °C, and subsequent transfer of the aqueous phase. To the sample, 0.1 volume of 3M sodium acetate was added and mixed, and 1 μL of GlycoBlue™ Coprecipitant (15 mg/mL) (Thermo Fisher Scientific) was included to aid in pellet visualization. To precipitate the DNA or RNA, 2.5-3 volumes of ice-cold ≥ 96% ethanol was added, followed by overnight incubation at −20 °C. Samples were then centrifuged at 15,200 rpm, 4 °C for 30-60 minutes, and the supernatant was discarded. Pellets were washed with 750 μL of ice-cold 70% ethanol, followed by centrifugation at 15,200 rpm, 4 °C for 15 minutes, and ethanol was removed. Pellets were left to dry for 10 minutes at room temperature and then re-suspended in nuclease-free water.

#### Ion pair-reverse phase high-performance liquid chromatography (IP-RP HPLC)

An UltiMate 3000 HPLC system (Thermo Fisher Scientific, UK) was utilized to perform IP-RP HPLC analysis with either a DNA Pack RP column (2.1 × 50 mm ID or 2.1 × 100 mm ID, Thermo Fisher) and UV detection at a wavelength of 260 nm. Weak IP-RP HPLC analysis was conducted under the following conditions: Buffer A, 100 mM triethylammonium acetate (TEAA) at pH 7.0, Buffer B, 0.1 M TEAA at pH 7.0 containing 25% acetonitrile. RNA samples were examined using different gradients, under native conditions (50 °C), starting at 40% buffer B to 45% in 1 minute, followed by a linear extension to 70% buffer B over 15 minutes, and then an extension to 90% buffer B over 2 minutes at a flow rate of 0.25 ml/min, and denaturing conditions (85 °C), starting at 35% buffer B to 45% in 1 minute, followed by a linear extension to 70% buffer B over 16 minutes, and then an extension to 90% buffer B over 2 minutes at a flow rate of 0.25 ml/min. The following ladders were used to determine size: peqGOLD 50b bp (VWR international), GeneRuler 1 kb Plus DNA Ladder (Thermo Fisher Scientific), or RiboRuler Low Range RNA ladder (Thermo Fisher Scientific).

#### Polymerase chain reaction

The Techne Primer thermocycler (Cole Parmer) was employed for polymerase chain reaction. KAPA2G Fast Hotstart Readymix (Merck) was used for all PCR reactions. Optimal cycling parameters for individual PCR reactions were determined following manufacturers’ guidelines. Purification of PCR products for downstream applications was accomplished using either phenol/chloroform ethanol precipitation or the Monarch® PCR & DNA Cleanup Kit (New England Biolabs) according to the manufacturer’s instructions.

#### Molecular cloning

Restriction enzymes were sourced from New England Biolabs or Fisher Scientific. Digestion parameters were followed according to the manufacturer’s instructions. GeneJet Gel extraction kit (Thermo Fisher Scientific) was used prior to ligation, following the manufacturer’s instructions. The Quick Ligation™ Kit (New England Biolabs) was used to ligate the vector and backbone fragments, following the manufacturer’s instructions. Competent cells were prepared using the Mix and Go! *E. coli* Transformation Kit (Zymo Research), and transformations were carried out using 50 ng of ligation product or plasmid, following the manufacturer’s instructions.

#### Bioassay

CPB larvae were fed on an artificial diet for the duration of the assay in a 48-well plate containing 500 µl diet per well. A range of equal numbers of *E.coli* cells expressing β-Actin dsRNA (NT vs 3T) was applied as an aqueous solution to the diet surface. The range of cells was generated via a 3–fold dilution series, performed 8 times. Plates were dried in a laminar flow hood. Per treatment, 24 larvae were screened, and mortality was scored on days 3, 4, 6, 7 and 10. An untreated control (no dsRNA) was included.

The artificial diet used, adapted from Gelman et al., (2001) (500 ml): - water (MilliQ) 384 mL, agar 7 g, rolled oats (ground) 20 g, torula yeast 30 g, lactalbumin hydrolysate 15 g, casein 5 g, fructose 10 g, wesson salt mixture 2 g, tomato fruit powder 6.25 g, potato leaf powder 12.5 g, β-sitosterol 500 mg, sorbic acid 400 mg, methyl paraben (nipagen) 400 mg, vanderzant Vitamin mix 6 g, neomycin sulfate 100 mg, aureomycin (chlortetracycline) 65 mg, rifampicin 65 mg, chloramphenicol 65 mg, nystatin 100 mg.

#### RNA characterisation by mass spectrometry

IP-RP HPLC-UV-MS analysis was performed on a Vanquish UHPLC system (Thermo Fisher Scientific) online to an Orbitrap Exploris 240 mass spectrometer (Thermo Fisher Scientific). Separations were performed using a 100 mm x 2.1 mm I.D. DNAPac RP column (Thermo Fisher Scientific). Mobile phase A was comprised of 10 mM dibutylamine (DBA) and 50 mM 1,1,1,3,3,3-hexaflouro-2-propanol (HFIP) whilst mobile phase B was comprised of 10 mM DBA and 50 mM HFIP with 50% acetonitrile. The HPLC gradient started at 35% B with a linear extension over 1 minute to 40% B. Mobile phase B was increased to 55% until minute 20 at a rate of ‘curve 3’ followed by a linear extension to 80% at 20.1 minutes for a 3-minute wash before returning to 35% at 23.2 minutes for a 10-minute equilibration stage and a total run time of 33.2 minutes. Separations were performed at a flow rate of 0.25 mL min^-1^ and a temperature of 50 °C with UV detection at 260 nm. The Orbitrap Exploris 240 was run using the Intact Protein application in Low Pressure mode. Spectra were captured at orbitrap resolutions of 15000 with a scan range of 450-2500 m/z, a normalised AGC target of 100%, 3 microscans, and 100 ms injection time. Samples were prepared with the addition of EDTA-free acid (adjusted to pH 8 with ammonium hydroxide) to a final concentration of 25 mM.

LC-MS data was analysed using BioPharma Finder 5.1 using the Intact Mass Analysis experiment type. 15000 Orbitrap resolution data was processed using the ReSpect™ deconvolution algorithm with the Sliding Windows Source Spectra Method. Sliding window parameters were adjusted to accommodate the chromatographic peaks with the Results Filters and Advanced Parameters set according to the potential RNA products from the expression constructs. The Peak Model was set to Nucleotide with a Negative Charge specified for deconvolution.

RNA secondary structure prediction was performed using MXfold2 (Sato et al., 2021) accessed through the Sato Lab web server.

The extinction coefficients of characterised RNA species were calculated using an online tool (available at molbiotools.com). Calculations employed the nearest neighbour method along with the characterised 260 nm molar extinction coefficient of the constituent nucleotides. The relative abundance of the terminated RNA products was calculated from the LC-UV chromatograms by considering the different extinction coefficients of the characterised RNA products and the chromatographic peak area. Integrated UV peaks for each of the characterised RNA products (mAU x min) were divided by their molar extinction coefficients (ε).

## Results and discussion

### Studying the effect of transcriptional terminators on the production of dsRNA in *E. coli*

#### Plasmid construct design

A series of plasmid constructs were designed for production of dsRNA in *E. coli* HT115(DE3) cells using two convergent T7 RNA polymerase promoters flanking a target dsRNA sequence (Dome 11, 400 bp), utilising the vector pMA-7 containing a ColEI origin of replication. A series of synthetic terminators were used based on those previously developed by Meirhofer et al., (2015) and Calvopina-Chavez et al., (2022). Primary work focused on the class I T7 synthetic terminator (T7 S) and the multiclass endogenous *E. coli* terminator, *rrnB* T1. From this we investigated four combinations of tandem class 1 and multiclass terminator sequences; T7 S, T7 S + *rrnB* T1, T7 S + *rrnB* T1 + T7 S, T7 S + *rrnB* T1 + T7 S + *rrnB* T1 flanking the target gene Dome11 (see Figure 1).

**Figure 1.**
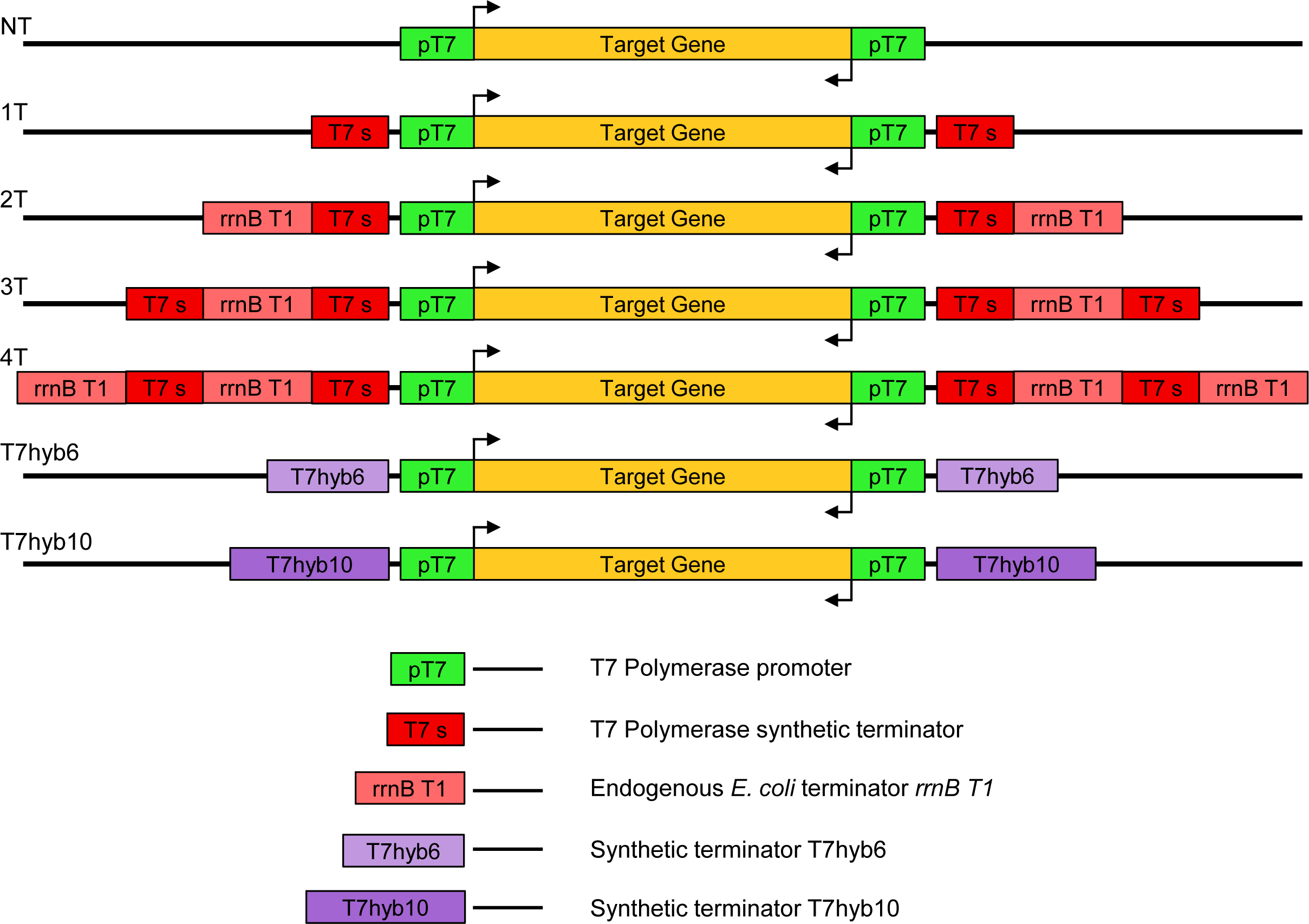
Schematic illustration of the plasmid constructs designed for dsRNA synthesis using transcriptional terminators. Schematics show the insert vectors utilised within the study not including the GeneArt pMA vector backbone which includes an ampicillin resistance gene and ColE1 origin of replication. Target genes included Dome11, β-Actin and GFP. The various T7 terminators and promoters used are indicated in the key.

Further work investigated two novel synthetic terminators developed by Calvopina-Chavez et al., (2022). The first, T7hyb6, consists of a class I terminator engineered to incorporate two slightly overlapping class II pause sites. The second, T7hyb10, is a tandem double hairpin terminator combining the novel synthetic terminators, T7hyb6 and T7hyb4, the latter a class I terminator, with a single pause site embedded in the poly-U-proximal segment of the terminator stem (see Figure 1). Terminator sequences can be found within Supplementary Table 1.

#### Cell growth - Dome11 dsRNA

Following the initial design and synthesis of the plasmid constructs described above, all plasmids expressing Dome 11 dsRNA including an additional control plasmid that does not contain the convergent dual T7 promoters (therefore unable to produce dsRNA) were transformed into *E. coli* HT115 cells. Three separate colonies were chosen as biological replicates and cultured in LB media overnight. Outgrowths and inductions were performed for each colony and growth curve measurements using OD_600_ nm readings were recorded every hour prior and up to 4 hours post-induction with ITPG (see Figure 2A). The results show a significant difference in cell growth post-induction (dsRNA synthesis) in those *E. coli* cells expressing dsRNA compared to the control (not expressing dsRNA). These results indicate that the maximum metabolic burden and toxicity within the cells is due to transcription and resulting formation of dsRNA in the cells and not the production of T7 RNA polymerase (consistent with previous observations (Delgado-Martin & Velasco, 2021)). The results also show that overall similar growth curves were obtained for *E. coli* cells transformed with each of the different plasmid constructs used in this study. Plasmids with the synthetic terminators (T7hyb6 and T7hyb10) had the highest OD_600_ post-induction, 1.54 and 1.48 respectively.

**Figure 2.**
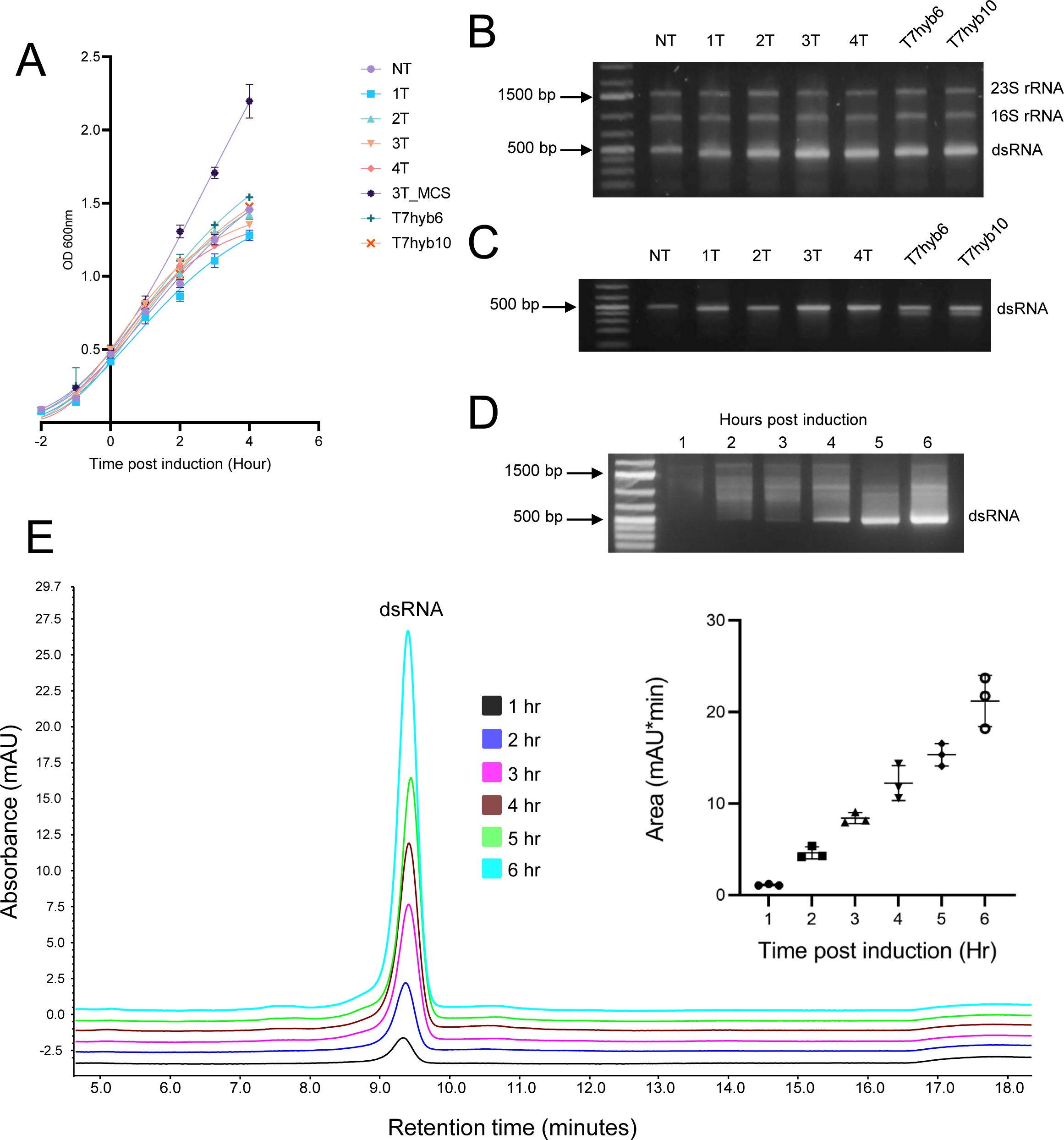
Investigations of transcriptional terminators on the production of Dome11 dsRNA. Plasmids pNT-4T Dome11, pT7hyb6/10 Dome11 and non-producing p3T_MCS were investigated **A.** Growth curve comparison of *E. coli* HT115 cells transformed with each of the plasmids. An outgrowth was performed and allowed to grow to an OD 600nm of 0.4-0.6. Samples were then induced for 4 hours, with OD measurements recorded at hour time points. Data shown as mean ± SD and are representative of three biological replicates. **B/C.** Agarose gel electrophoresis analysis of RNA extractions in the absence (**B**) and presence of RNase T1 (**C**). The corresponding rRNA and Dome 11 dsRNA (400 bp) are highlighted. **D/E** Analysis of dsRNA extracted from p3T_Dome11 HT115 cells at different time points post induction**. D** Agarose gel electrophoresis analysis. **E.** IP-RP HPLC chromatography analysis. Inset shows the relative peak area (mAU*min), measured using IP-RP HPLC chromatography. Data is shown as mean ± SD and are representative of 3 biological replicates.

#### Quantitative analysis of dsRNA production in *E. coli* - Dome11

For RNA analysis, aliquots of 1 x 10^9^ cells (4 hours post induction) were used prior to RNA extraction and purification in both the absence and presence of RNase T1 to remove the *E. coli* ssRNA (tRNA and rRNA). Agarose gel electrophoresis analysis of the extracted RNA is shown in Figure 2B. The results show the successful synthesis of dsRNA Dome11 (_≅_ 400 bp) in each of the different plasmid designs. Semi-quantitative analysis based on gel band intensity shows that the p3T_Dome11 and p4T_Dome11 resulted in the highest dsRNA yields. In addition, RNA extractions were performed in the presence of RNase T1 confirming the dsRNA product (resistant to RNase T1) and the semi-quantitative analysis again highlights the increased dsRNA yield as the number of transcriptional terminators increases (see Figure 2B).

A further study was performed to investigate the production of dsRNA at various time points post induction (1-6 hours) in *E. coli.* Three p3T_Dome11 colonies were selected, grown, and induced as previously described, however, the induction time was extended to 6 hours. For RNA analysis, RNA was extracted from 1 x 10^9^ cells prior to agarose gel electrophoresis and ion pair reverse phase HPLC (IP-RP-HPLC). Agarose gel electrophoresis analysis indicates that the yield of dsRNA increases up to 6 hours post-induction (Figure 2D). This is further confirmed via relative quantification using IP RP HPLC (Figure 2E). At 6 hours post-induction, a mean relative yield value of 21.12 was determined, compared to 1-hour post-induction, with a mean relative yield of 1.13. These results confirm that total dsRNA yield increases with hours post induction up to 6 hours and potentially longer during small-scale batch fermentation.

Accurate quantification of dsRNA yield was performed using both relative quantification of the dsRNA in conjunction with IP RP HPLC analysis and absolute quantification of the dsRNA using UV spectrophotometry following purification of the dsRNA. IP-RP-HPLC analysis of the extracted dsRNA is shown in Figure 3A. The results show the presence of the endogenous rRNA and dsRNA synthesised in *E. coli* from each of the plasmid constructs used in this study. Relative dsRNA quantification was performed using the peak areas from the resulting chromatograms (Figure 3A and summarised in Figure 3B). The results show that consistent with the semi-quantitative gel electrophoresis analysis as the number of transcriptional terminators present in the plasmid construct increases (0-3), the amount of dsRNA produced also increases. The dsRNA produced from the plasmid construct with either three or four transcriptional terminators (p3T_Dome11 and p4T_Dome11) produced the highest relative yields of dsRNA respectively (17.26 and 17.29), with no significant difference between the two. Alternative hybrid terminators also demonstrated an increase in the amount of dsRNA produced compared to no terminators, with pT7hyb6 and pT7hyb10 producing relative yields of 10.49 and 14.37, respectively. These results indicate that pT7hyb6 produces a significantly similar amount of dsRNA to p2T_Dome11 (10.12). Comparing the relative amount of dsRNA produced in the absence of transcriptional terminators (pNT_D11 =2.219), to dsRNA produced using 3 transcriptional terminators (p3T_D11 =17.26), the results show an increase of 7.78-fold.

**Figure 3.**
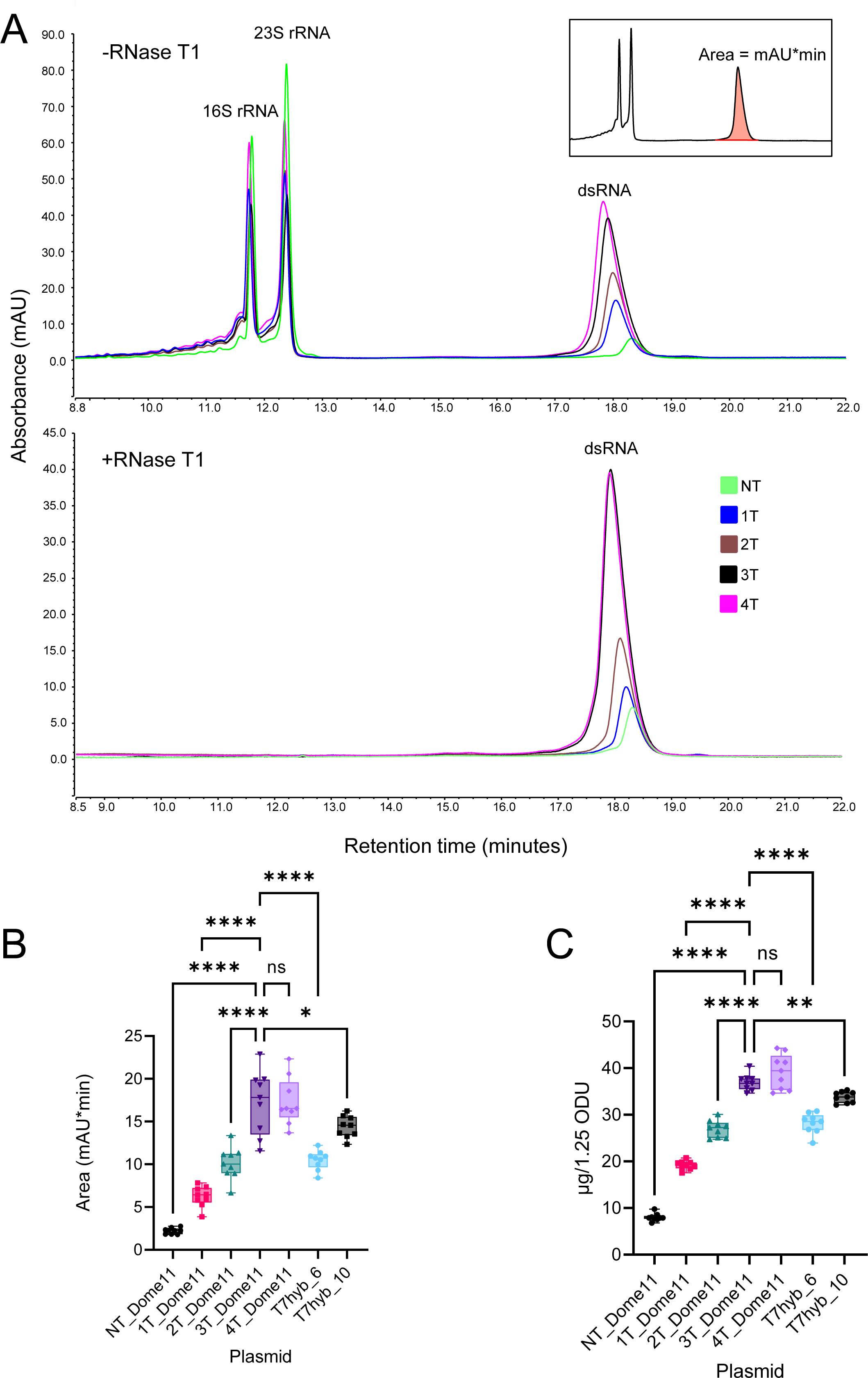
Quantification of dsRNA yield. **A**. IP-RP HPLC chromatograms of RNA extracts from *E.coli* HT115 cells transformed with pNT-4T_Dome11 and pT7hy6/10_Dome11. RNA extractions were performed in the absence and presence of RNase T1. The corresponding dsRNA and rRNA are highlighted. **B.** RNA samples were analysed using relative peak area (mAU*min). Significance was calculated using a one-way ANOVA test against p3T_Dome11. p3T_Dome11 and p4T_Dome11 demonstrate the highest relative yield of, 17.26 and 17.29, respectively, with no significant difference. Data represents triplicate technical replicates of three biological replicates; p<0.05 = (*), p<0.01 = (**), p<0.001 = (***), p<0.0001 (****). **C.** Comparison of absolute dsRNA yield of pNT-4T_Dome11 and T7hyb6/10_Dome11 RNA samples. RNA samples were analysed using UV spectrophotometry to determine RNA concentration using a mass concentration/A260 unit (46.52 μg/mL). Significance was calculated using a one-way ANOVA test against p3T_Dome11. p3T_Dome11 and 4T_Dome11 demonstrate the highest absolute yield of, 36.95 µg and 39.22 µg, respectively, with no significant difference. Data is shown as box plots of triplicate technical replicates and are representative of three biological replicates; p<0.05 = (*), p<0.01 = (**), p<0.001 = (***), p<0.0001 (****).

Absolute quantification of the dsRNA produced in *E. coil* was also performed using a modified RNA extraction protocol including the addition of RNase T1 to remove any ssRNA from the extracted lysate prior to purification. IP-RP-HPLC analysis of the extracted dsRNA in the presence of RNase T1 is shown in Figure 3A and demonstrates the effective removal of rRNA leaving on the dsRNA present. Following purification, RNA samples were analysed using a UV spectrophotometry to determine RNA concentration using a mass concentration/A_260_ unit (46.52 μg/mL) as previously determined for dsRNA (Nwokeoji et al., 2017). The absolute quantification results are summarised in Figure 3C and show a consistent pattern as previously determined for the relative dsRNA quantification from the different transcriptional terminators used. Total dsRNA analysis indicated that using three or more transcriptional terminators (p3T_Dome11 and p4T_Dome11) produced the highest total yields of dsRNA, 36.95 µg and 39.22 µg respectively (from 1 x 10^9^ cells). Alternative hybrid terminators also demonstrated an increase in total dsRNA produced compared to the absence of transcriptional terminators, with pT7hyb6 and pT7hyb10 producing yields of 28.19 µg and 33.78 µg, respectively. These results indicate that pT7hyb6 produces a significantly similar amount of total dsRNA to p2T_Dome11, 26.95 µg. Comparing the total dsRNA produced without the use of transcriptional terminators-pNT_Dome11 (8.028 µg), to dsRNA with three transcriptional terminators, p3T_Dome11 (36.95 µg), the results show a 4.94-fold increase.

In summary, the results show that utilising multiple synthetic transcriptional terminators outside of the dual convergent T7 RNA polymerase promoters in the plasmid construct design results in a significant increase in the amount of dsRNA biocontrol expressed *in vivo* in *E. coli* HT115 cells.

#### High yield production of dsRNA in a bioreactor

Following initial work in small-scale shake flasks (250 ml flasks /50 ml cultures) further work was performed comparing dsRNA production in industry-relevant bioreactors. A 1 L bioreactor with a working volume of 0.75 L was used. *E. coli* HT115 cells transformed with either (pNT_Dome11, p1-3T_Dome11) were grown overnight in a shake flask prior to a 5% (v/v) inoculum in the bioreactor. Following induction with IPTG (1 mM) samples were aseptically withdrawn from the bioreactor and OD_600_ nm was taken (see Figure 4A). The results show that the highest cell density (prior to cell death) was obtained by p3T_Dome 11 in comparison to the constructs with no terminators or a single terminator. During the course of the fermentation experiment 1 x 10^9^ cells were taken prior to RNA extraction and quantification using IP RP HPLC (see Figure 4B/C). The results show an increase in both the relative and absolute amounts of dsRNA produced at multiple time points during the fermentation using the triple terminator dsRNA in comparison to no terminators. Comparing the total dsRNA produced from 1 x 10^9^ cells (1.25 AU) at the 6.5 hours post induction time point without the use of transcriptional terminators pNT_Dome11 (21 µg), to dsRNA with three transcriptional terminators p3T_Dome11 (116.66 µg) the results show a 5.56-fold increase. These results are consistent with the previous small-scale shake flask experiments and demonstrate that increased dsRNA yield is achieved using triple terminators compared to no terminators. Furthermore, combined with the data demonstrating that increased cell density was achieved during the fermentation for the p3T_Dome 11, this results in a significantly higher overall yield of dsRNA from the large-scale fermentation.

**Figure 4.**
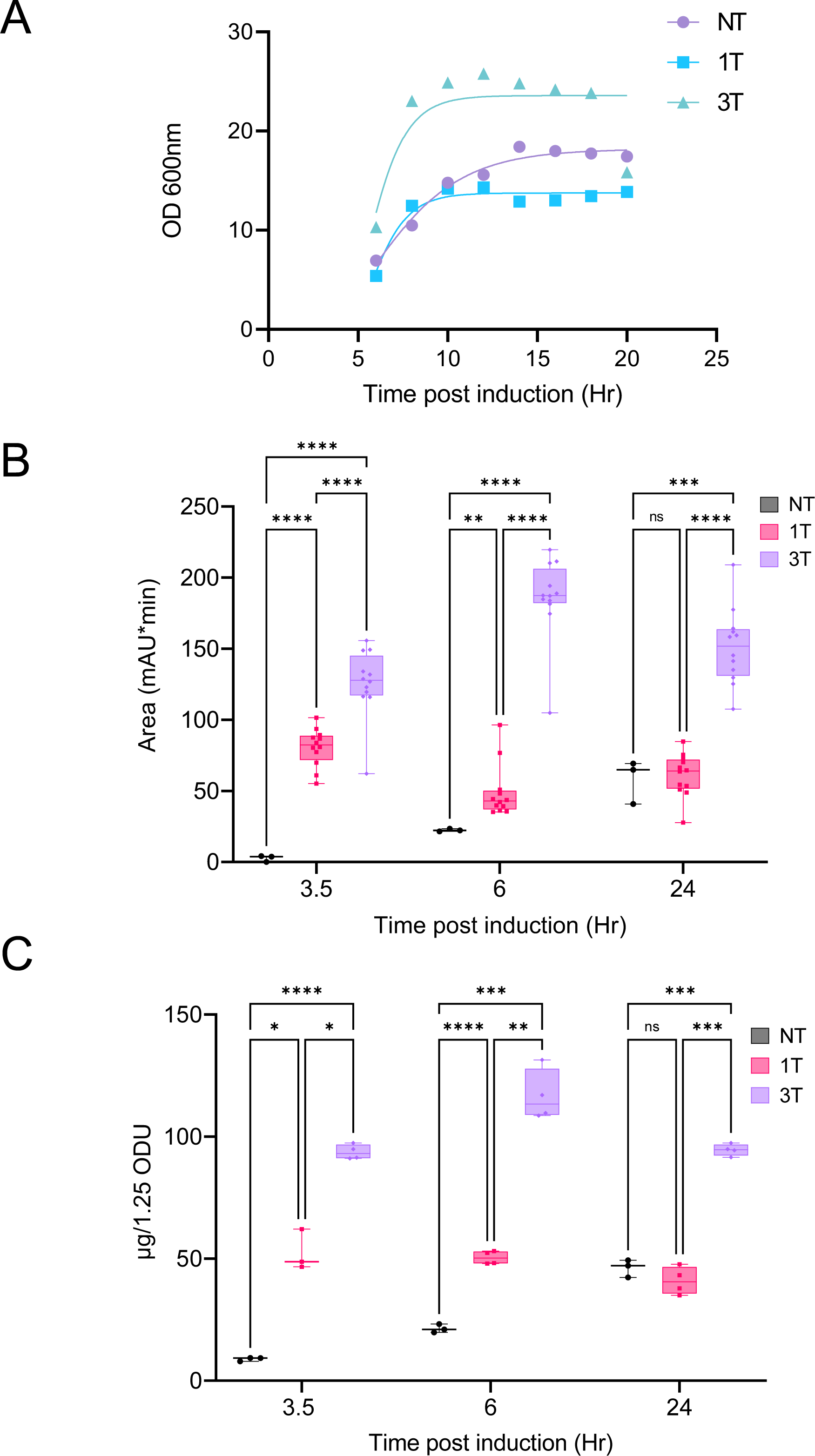
Large-scale batch fermentation production of Dome11 dsRNA. Plasmids pNT/1T/2T/3T Dome11 were investigated **A.** Growth curve comparison of *E. coli* HT115 cells transformed with each of the plasmids. Data shown is representative of a single biological replicate. **B.** RNA samples were taken 3.5, 6 and 24 hours post induction and analysed using relative peak area (mAU*min). Significance was calculated using Tukey’s multiple comparisons test against pNT_Dome11 (N=3), p1T_Dome11 (N=12) and p3T_Dome11 (N=12). p3T_Dome11 demonstrates the highest relative yield of 185.711 at 6.5 hours post induction. p<0.01 = (**), p<0.001 = (***), p<0.0001 (****). **C.** Comparison of absolute dsRNA yield. RNA samples were analysed using a UV spectrophotometry to determine RNA concentration using a mass concentration/A260 unit (46.52 μg/mL). Significance was calculated using Tukey’s multiple comparisons test against pNT_Dome11 (N=3), p1T_Dome11 (N=4) and p3T_Dome11 (N=4), an outlier was removed at for p1T_Dome11 at time point 3.5 hours post-induction. p3T_Dome11 demonstrates the highest absolute yield of 116.66 µg at time point 6.5 hours post-induction. p<0.05 = (*), p<0.01 = (**), p<0.001 = (***), p<0.0001 (****).

Both the dsRNA/biomass yield and maximum productivity were determined from the p3T_Dome 11 large scale fermentation following methods by Papic et al., (2018). A dsRNA/biomass yield of 0.065 g·g^-1^ and a maximum productivity of 34.3 mg l^-1^ h^-1^ at 3.5 hours post-induction was obtained. These results demonstrate a 2.25-fold increase in productivity compared to the maximum productivity obtained using previous fed-batch experiments, 15.2 mg l^-1^ h^-1^ (Papic et al., 2018). To summarise our data demonstrates that batch fermentation production dsRNA using multiple transcriptional terminators is scalable and generates significantly higher yields of dsRNA generated in the absence of transcriptional terminators from both small-scale batch and large-scale fermentation conditions.

#### Quantitative analysis of dsRNA production in *E. coli* - β-Actin and GFP dsRNA

Having demonstrated the increased yield of dsRNA (Dome11 - 400 bp dsRNA) using multipleT7 transcriptional terminators, further studies were performed to analyse the effect of transcriptional terminators on the production of two alternative dsRNA sequences (β-Actin and GFP). Five more plasmid constructs utilising dual convergent T7 promoters flanking different target genes were designed, two for β-Actin (NT and 3T) and three for GFP (NT, 1T and 3T). Each plasmid construct was then transformed into *E. coli* HT115 cells, prior to a series of small-scale shake flask inductions and RNA extraction as previously described. Three biological replicates for each analysis and the resulting cell growth curves are shown in Figure 5A. The results show that *E. coli* cells expressing dsRNA increase the metabolic burden induced onto the cells which is consistent with previous data. Agarose gel electrophoresis of the dsRNA extracted for β-Actin and GFP (302 bp and 263 bp respectively) in the presence and absence of RNase T1 are shown in Figure 5B-E. Semi-quantitative analysis based on gel band intensity shows an increase in the amount of dsRNA as the number of transcriptional terminators is increased, consistent with results obtained for Dome 11 dsRNA.

**Figure 5.**
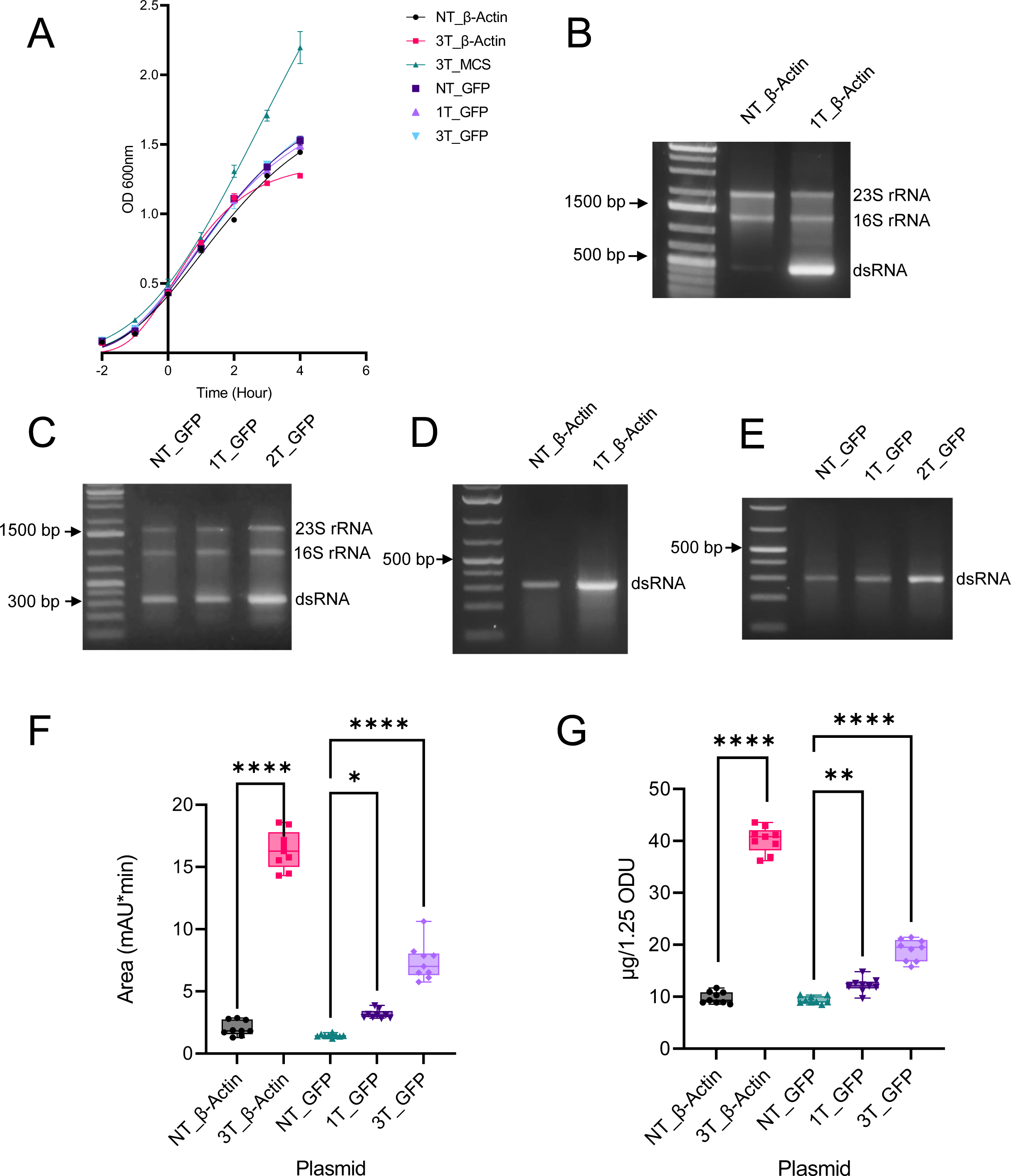
Investigations of transcriptional terminators on the production of β-Actin and GFP dsRNA. **A.** Growth curve comparisons of *E. coli* HT115 cells transformed pNT/3T_β-Actin and pNT/1T/3T_GFP and the control non dsRNA producing p3T_MCS. Data shown as mean ± SD and are representative of three biological replicates **B/C/D/E.** Agarose gel electrophoresis analysis of RNA extractions in the absence (B) and presence of RNase T1 (C). The corresponding rRNA and Dome 11 dsRNA (400 bp) are highlighted. The corresponding dsRNAs are highlighted (302 bp and 263 bp) is noted for in all constructs. **F.** Comparison of relative dsRNA yield of pNT/3T_β-Actin and pNT/1T/3T_GFP RNA samples. RNA samples were analysed using relative peak area (mAU*min), measured via IP-RP HPLC chromatography. Data represents triplicate technical replicates of three biological replicates. Independent statistical tests were performed on β-Actin (unpaired T-test) and GFP samples (Kruskal-Wallis), p<0.05 = (*), p<0.01 = (**), p<0.001 = (***), p<0.0001 (****). p3T_ β-Actin and p3T_GFP demonstrated the highest relative yield, 16.36 and 7.385, respectively. **G.** Comparison of absolute dsRNA yield of pNT/3T_β-Actin and NT/1T/3T_GFP RNA samples. RNA samples were analysed using a UV spectrophotometry to determine RNA concentration using a mass concentration/A260 unit (46.52 μg/mL). Data represents triplicate technical replicates of three biological replicates. Independent statistical tests were performed on β-Actin (unpaired T-test) and GFP samples (one-way ANOVA), p<0.05 = (*), p<0.01 = (**), p<0.001 = (***), p<0.0001 (****). p3T_ β-Actin and p3T_GFP demonstrated the absolute dsRNA, 40.26 µg and 19.08 µg, respectively.

Further quantitative analysis of the relative yields of β-Actin and GFP dsRNA was performed by IP-RP-HPLC (see Supplementary Figure 1). Relative and absolute quantification was performed as previously described and the results are summarised in Figure 5F/G. The results show an 8-fold increase in the relative abundance of β-Actin dsRNA using the triple transcriptional terminators in comparison to the dsRNA generated without transcriptional terminators. The relative abundance of the GFP dsRNA also increased 5-fold when comparing the yield of dsRNA with triple transcriptional terminators in comparison to the dsRNA generated without transcriptional terminators.

Absolute quantification of the dsRNA was also performed as previously described. The dsRNA analysis of β-Actin indicated that p3T_β-Actin produced the highest total yields of dsRNA (40.26 µg). Comparing the total dsRNA yield from 1 x 10^9^ cells in the absence of transcriptional terminators (pNT_β-Actin = 9.967 µg), to the use of three transcriptional terminators (p3T_β-Actin = 40.26 µg) demonstrates a 4-fold increase. Total dsRNA analysis of GFP indicated that p3T_GFP produced the highest total yields of dsRNA (19.08 µg). Comparing the total dsRNA yield of non-terminated dsRNA produced by pNT_GFP (9.297 µg), to p3T_GFP (19.08 µg) with a fold increase of 2.05. It is interesting to note that there is variation in yields of dsRNA produced by all the triple terminator constructs, 36.95 µg (Dome11), 40.26 µg (β-Actin), and 19.08 µg (GFP). These results demonstrate that the production of target dsRNA is likely dependent on the sequence and size of the corresponding dsRNA.

In summary, we have demonstrated that the addition of multiple transcriptional terminators outside of the dual convergent T7 promoters leads to a significant increase in the production of a range of dsRNAs of different sequences and sizes. Therefore, these results demonstrate the versatility of this approach that can be potentially used for production of a wide range of different dsRNA biocontrols across multiple gene targets and can be implemented into other designs if necessary.

#### Increased bioactivity of dsRNA using multiple transcriptional terminators

Having demonstrated the increase in dsRNA yield generated from plasmid constructs with multiple transcriptional terminators, further work was performed to demonstrate an associated increase in RNAi efficacy when used as a dsRNA biocontrol. Following cell growth and induction of *E. coli* HT115 cells expressing either β*-Actin* dsRNA (NT vs 3T), insect-feeding assays were set up and performed on L2 larvae of the Colorado potato beetle. An equal number of *E. coli* cells expressing β*-Actin* dsRNA (NT vs 3T) were taken and a 3–fold dilution series was performed 8 times. A total of 24 insects were studied per dilution across both treatments. Insect mortality percentage was scored across 10 days and a negative control was used (-dsRNA) to indicate baseline mortality levels. Two biological replicates were performed and the results are shown in Figure 6.

**Figure 6.**
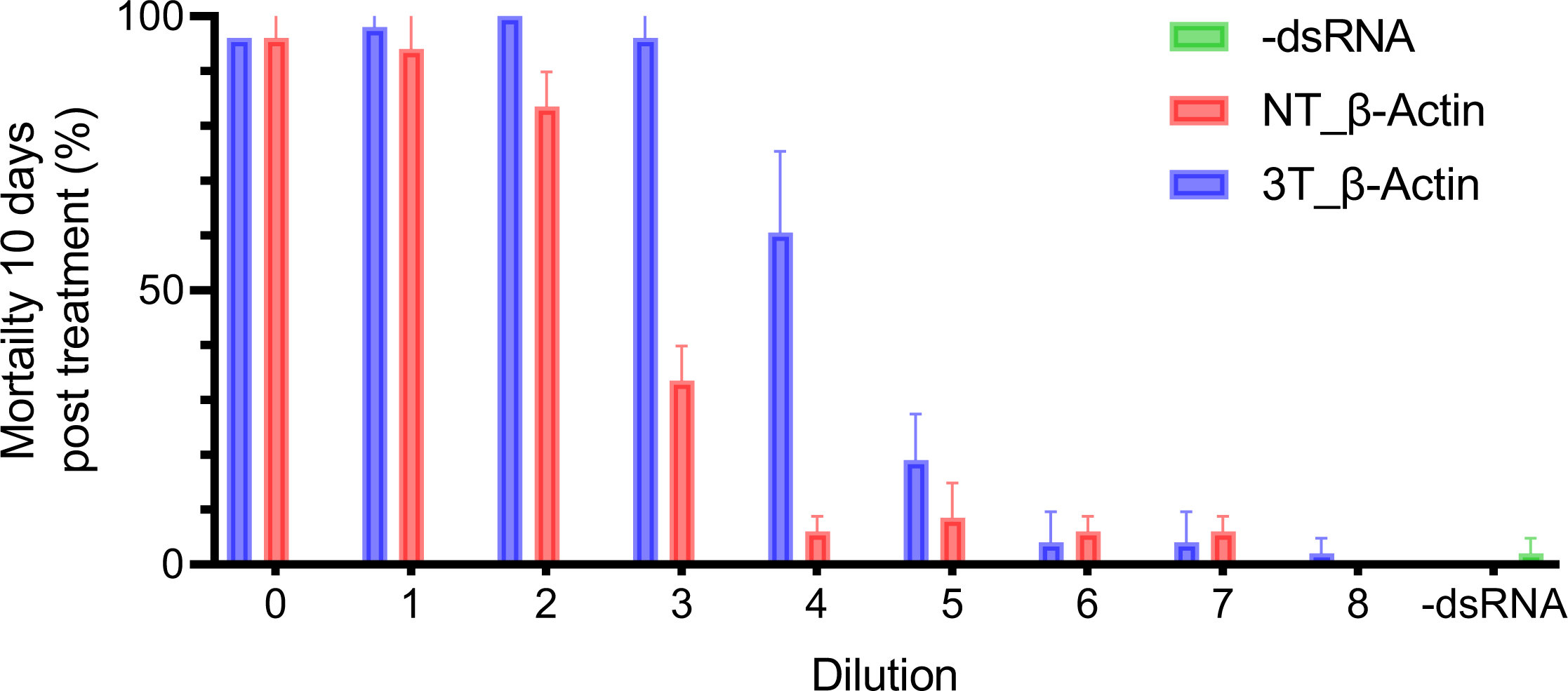
*In vivo* RNAi efficacy of dsRNA biocontrols produced from DNA constructs in the presence and absence of transcriptional terminators. An insect feeding assay was performed to investigate the effect of equal amounts of dsRNA *E.coli* cells expressing either un-terminated dsRNA (NT_β-Actin) and terminated dsRNA (3T_β-Actin). A series of 8 dilutions of *E.coli cells* and two independent assays were performed. Mortality percentage was scored across 10 days. A negative control was used (-dsRNA) to indicate baseline mortality levels. Data shown as mean ± SD of two biological replicates.

The results show that for the first two dilutions similar levels of average mortality (%), were observed for the *E. coli* cells with NT and 3T β*-Actin* dsRNA. However, comparing the third dilution, the average mortality (%) observed for L2 larvae fed with *E. coli* cells expressing NT β*-Actin* dsRNA was 33.5% compared to 96% for the 3T β*-Actin* dsRNA. This trend was also observed in the following dilution where 6% and 60.5% average mortality was determined for the NT and 3T β*-Actin* dsRNA.

These results demonstrate that both the NT and 3T β*-Actin* dsRNA successfully act as dsRNA biocontrols *in vivo* as demonstrated by live insect feeding assays using L2 larvae of the Colorado potato beetle. Moreover, the results show that using the same number of *E. coli* cells expressing 3T β*-Actin* dsRNA results in higher RNAi efficacy as measured by insect mortality compared to *E. coli* cells expressing NT-dsRNA. These results are consistent with previous quantitative analysis described above that demonstrated a 4.93-fold increase in the total dsRNA yield of the 3T-dsRNA compared to NT-dsRNA, 15.6 ug/1×10^9^ cell and 76.9 ug/1×10^9^ cell, respectively. This data indicates that using microbial cells expressing dsRNA using transcriptional terminators increases the RNAi efficacy of dsRNA biocontrols.

#### Measuring transcriptional termination efficiency

A series of *in vitro* transcription (IVT) assays were performed to investigate the transcriptional termination efficiency of the different transcriptional terminators in the dual convergent T7 RNA polymerase promoter plasmid construct designs. Plasmids with NT – 4T expressing the *Dome11* dsRNA were linearised to produce a single cut in the AmpR site. Different-sized RNA fragments will be generated upon either termination at the transcriptional terminators or run-off transcription at the end of the linearised DNA template (see Supplementary Figure S2). Gel electrophoresis analysis of the dsRNA synthesised using IVT from these linearised DNA templates is shown in Figure 7A.

**Figure 7.**
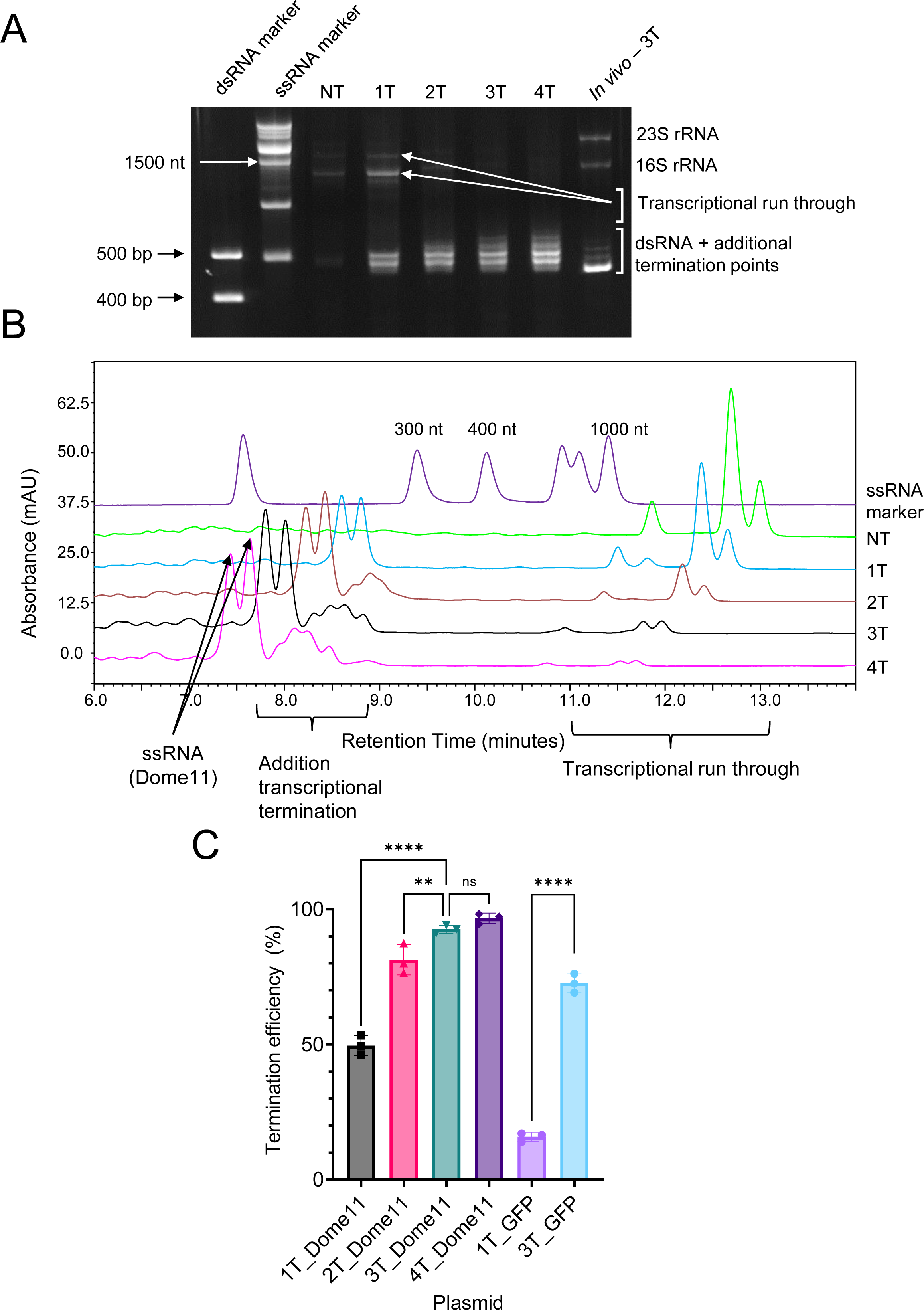
Measuring transcriptional termination efficiency *in vitro*. **A**. Ex-Gel electrophoresis analysis of in vitro transcription products. An *in-vivo* produced Dome11 dsRNA was also run as a comparison. **B.** IP-RP HPLC chromatogram of in vitro transcription products analysed under denaturing conditions, 85 °C, using the gradient 45 - 50 - 66, 15 mins. Marker = RiboRuler Low Range RNA Ladder. Samples offset by signal (5%) to allow for clear visual comparison and indicate products < 1000 > (nts). **C.** Comparison of the transcriptional termination efficiency between production systems pNT/1T/2T/3T/4T Dome11 and p1T/3T GFP. Data shown as mean ± SD of three biological replicates. Independent statistical tests were performed on Dome11 (one-way ANOVA) and GFP samples (unpaired T-test), p<0.05 = (*), p<0.01 = (**), p<0.001 = (***), p<0.0001 (****).

The results show that the addition of multiple terminators (NT to 4T), results in a reduction in ssRNA transcriptional run-through products (expected ssRNA products of 1381 to 1562 nts or 1873 to 2632 nts as shown on Supplementary Figure S2 in conjunction with an increase in the amount of the terminated RNAs of the expected size range (expected ssRNA 427 – 581 nts, depending on the termination point). The samples were run alongside an *in vivo Dome11* dsRNA produced and extracted from *E. coli* HT115 cells as a control, to allow a comparison of which band indicates the *Dome11* dsRNA with no ssRNA overhangs (see Figure 7A). It is interesting to note that when comparing the *in vitro* and *in vivo* generated dsRNA, the *in vitro* dsRNA products result in a range of additional dsRNA species of increasing size in contrast to dsRNA produced and extracted *E. coli* HT115 where only minor low abundance species are observed. It is proposed that the difference in specific RNA populations produced *in vitro* vs *in vivo* is reflected in the different production and extraction conditions. *In vivo* produced dsRNAs (+ ssRNA overhangs) are subjected to endogenous RNases present in the *E. coli* and therefore the majority of the final products will have the ssRNA overhangs removed (see Supplementary Figure S3). In contrast, *in vitro-*produced RNA should be unaffected by such issues, therefore most of the products will be dsRNA with ssRNA overhangs resulting from potentially different termination points.

Following initial agarose gel electrophoresis analysis, further quantitative analysis was performed using IP-RP-HPLC under denaturing conditions (85 °C) to analyse the resulting ssRNAs to determine the transcriptional termination efficiency based on the amount of transcriptional run-through or successful transcriptional termination. The results are shown in Figure 7B and demonstrate that upon the addition of transcriptional terminators the amount of transcriptional run-through is decreased in conjunction with a resulting increase in ssRNA Dome11 (+ downstream termination). Quantification was measured based in the relative peak area of all the transcriptional run-through products and the terminated ssRNA products resulting in transcriptional termination efficiency of 92.65% (3T) and 97.75% (4T) compared to (49.72%) for a single terminator (1T) (Figure 7C).

Experiments were repeated with pNT/1T/3T_GFP expressing the *GFP* dsRNA. Results demonstrated transcriptional termination efficiency of 72.64% for triple terminators (3T) and 15.87% for a single terminator (1T). Considering potential upstream and downstream sequence effects that have been observed to impact the termination efficiency of T7 RNA polymerase, these results are consistent with a previous study where transcriptional terminators were utilised in alternative systems for the production of ssRNA where >90% termination efficiency was achieved using multiple transcriptional terminators (26). The differences illustrated in termination efficiency between target sequences consequently could signify the differences noticed in both relative and total *in vivo* yields.

#### Identification of sites of transcriptional termination using mass spectrometry analysis

Previous work described above showed that using multiple transcriptional terminators (2T-4T) resulted in increased transcriptional termination efficiency when measured *in vitro*. However, multiple products were observed when *in vitro* synthesised dsRNA was analysed using both agarose gel electrophoresis and denaturing IP-RP HPLC due to transcriptional termination potentially at multiple points across the multiple terminators. Therefore, further analysis was performed to provide further mechanistic insight into the transcriptional termination by identifying the different RNA species generated by using multiple transcriptional terminators in conjunction with mass spectrometry analysis. For mass spectrometry studies, we designed model systems to generate either a 50 nt ssRNA or 120 bp dsRNA from constructs containing the three terminator sequences (T7 S + *rrnB* T1 + T7 S). Due to low ionisation efficiency and associated metal ion adduction, large RNA molecules are not amenable to mass spectrometry analysis therefore shorter RNA lengths more amenable to intact mass spectrometry analysis were utilised in this study.

#### RNA characterisation using mass spectrometry analysis

Short-model RNAs were produced through *in vitro* transcription to study the composition of the terminated RNA products using intact mass analysis. Chromatograms from the 50 nt ssRNA construct show the presence of at least four major RNA species, supporting the prediction that the multiple products seen in agarose gels and IP-RP HPLC analysis of the dsRNAs correspond to transcriptional termination occurring at multiple points across the different transcriptional terminators (Figure 8A). Intact mass analysis of these species reveals four major RNA products differing in their 3’ ends due to alternative termination positions (Figure 8A). The most abundant species produced by the T7 RNA polymerase was found to correspond to transcription terminating at the end of the first T7 S terminator sequence (see Figure 8B), the next two most abundant species were found to correspond to transcriptional termination at different positions within the larger *rrnB* T1 terminator (Figure 8C/D). The final species characterised through mass spectrometry was found to be terminated once again at the end of the T7 S, the third and final terminator within the construct (Figure 8E). Due to low signal intensity and poor signal-to-noise ratio, it has not been possible to deconvolute intact masses for the more minor peaks such as those at 9.2 and 10.1 minutes. It is expected that these will constitute alternative sites of transcriptional termination.

**Figure 8.**
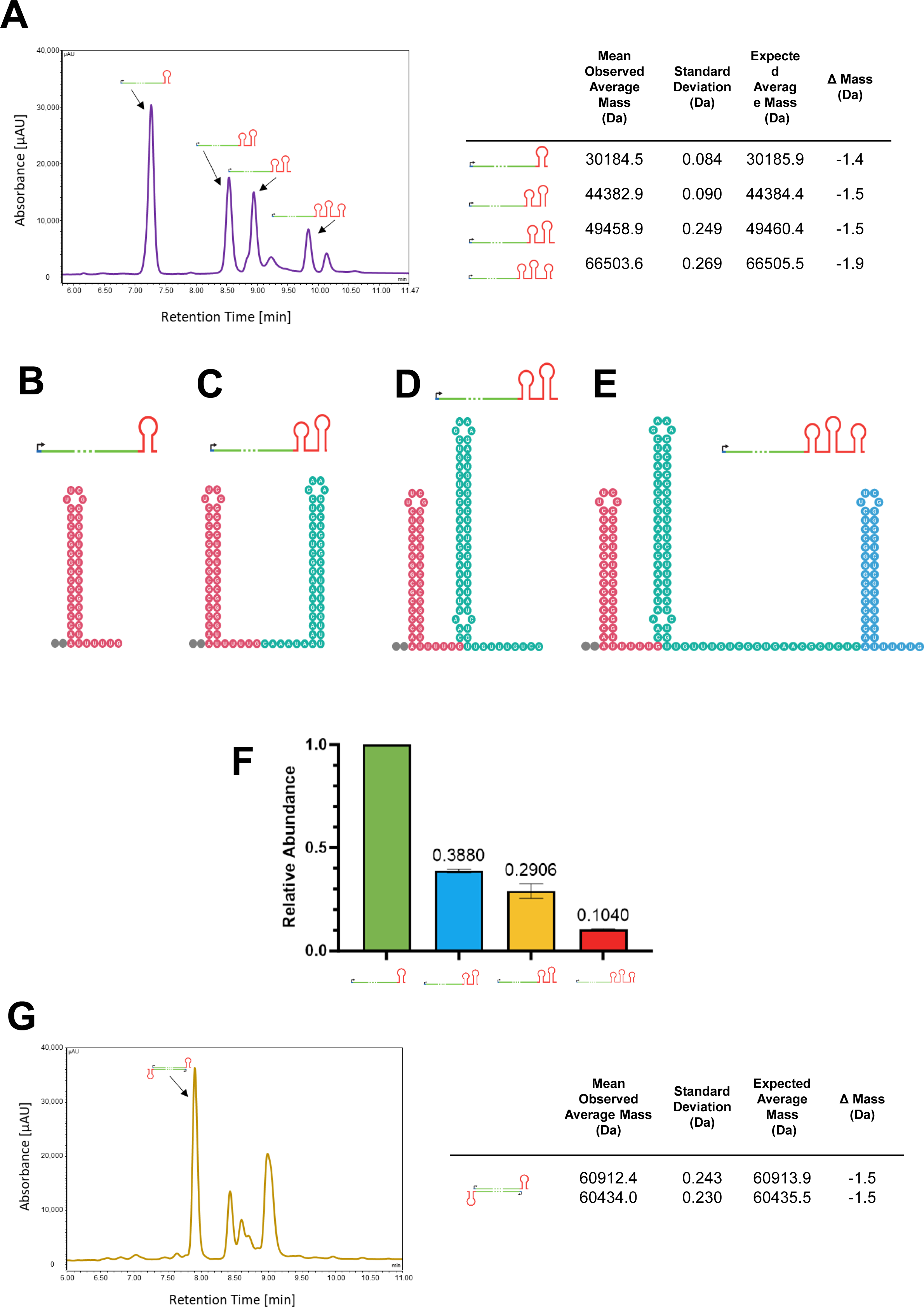
Characterisation of transcriptional termination using mass spectrometry. **A**. LC-UV chromatogram of the ssRNA *in vitro* transcribed from a linear DNA construct containing three transcriptional terminators (T7 S + *rrnB* T1 + T7 S). Deconvoluted average masses for each of the RNA species is highlighted and the corresponding RNA products determined from the mass spectrometry analysis are annotated with cartoons of RNA transcripts with differing numbers of stem-loop structures to represent the different terminator sequences present in the transcribed RNA. **B-E**. Predicted secondary structures of the transcribed terminator sequences. Minimum free energy secondary structure elements from the multiple termination positions within the triple terminator construct. Cartoons illustrate the different termination sites as different stem loops for each terminator sequence. Sequence corresponding to the different terminators is shown in different colours: First (T7 S), pink; second (*rrnB T1*), green; third (T7 S), blue. **F.** Relative abundance of the identified RNA products resulting from transcriptional termination. Mean relative abundances are plotted compared to the most intense product peak with error bars showing standard deviation (N=3). **G**. LC UV chromatogram of dsRNA *in vitro* transcribed from a linear DNA construct containing three transcriptional terminators (T7 S + *rrnB* T1 + T7 S). Deconvoluted average masses for each of the RNA species are shown with cartoons of RNA transcripts with differing numbers of stem-loop structures to represent the different terminator sequences present in the transcribed RNA.

Examination of the predicted minimum free energy RNA secondary structures of these terminator structures provides insight into the mechanistic basis of transcriptional termination when these terminator sequences are combined in an RNA expression construct (Figure 8B-E). The predicted structure of the terminator region of the major RNA product shows the canonical stem-loop secondary structure associated with intrinsic transcriptional terminators of a G-C rich stem, a short loop, and a U-rich tail (Figure 8B). The mass spectrometry analysis shows that transcription terminates after the run of four uracils and a single guanine in agreement with the termination position reported for the wild-type TΦ T7 terminator on which the T7 S terminator is based (Dunn & Studier, 1983). The next most abundant products contain sequences from the T7 S and *rrnB* T1 terminators. The predicted secondary structures suggest the presence of two stem-loop structures and the mass spectrometry analysis predicts two different sites of termination within the *rrnB* T1 sequence (Figures 8C/D). In the first of these two products, transcription terminates immediately after a predicted stem-loop structure following three uracils that constitute the end of the hairpin structure (Figure 8C). The second and lower abundance *rrnB* T1 termination product is predicted as having a larger stem-loop structure for the second terminator with termination occurring shortly afterwards with a run of nucleotides ending in UCG (Figure 8D). The second of these termination positions is consistent with the termination position reported by Song and Kang downstream of the conserved PTH sequence region (ATCTGTT) (Song & Kang, 2001). Consistent with the identification of potential upstream termination from the PTH sequence, we also identify another termination position upstream of the PTH sequence (Lubkowska et al., 2011) however identify cleavage after the U immediately prior to the conserved motif. We hypothesise that this different termination position is a result of the upstream sequence context of the *rrnB* T1 caused by the T7 S terminator.

Numerous studies have reported the importance of the overall sequence context and influence of this upstream region termination efficiency with the potential for competing RNA structures (Cambray et al., 2013; Carothers et al., 2011; Larson et al., 2008; Lubkowska et al., 2011; Yanofsky, 2000; Yoo & Kang, 1996). It has also been reported that the hairpins with a longer stem or stabilised loop can nucleate before the elongation complex reaches the pause point at the U-tract position (Penno et al., 2015). For transcription terminating at the end of the third and final terminator, the prediction shows three stem-loop structures with the structure of the T7 S terminator at the end of a run from the remainder of the *rrnB* T1 terminator sequence (Figure 8E). Transcription terminates at the same run of four Us and a G, similar to the first termination position.

Having characterised the identity of the major RNA products, their relative abundance was considered to measure the effect of termination efficiency at the different identified termination positions. Calculated extinction coefficients were used in conjunction with the chromatographic peak areas to determine the relative abundance for each of the products accounting for any differences in UV absorbance (see Figure 8F). The results show the majority of the termination occurs at the end of the T7 S terminator (the first terminator) with lower but additional termination occurring at the downstream terminators.

To verify that the transcriptional termination is consistent in a two convergent T7 promoter system, dsRNA was produced from the 120 bp expression construct. Once again multiple peaks were seen in the chromatograms, as with the 50nt ssRNA construct. Mass spectrometry analysis revealed that the major product was formed from sense and antisense strands terminating at the end of the first T7 S terminator sequence (Figure 8G). The other peaks observed in the chromatogram are believed to correspond to the other termination positions as seen with the 50nt ssRNA. It has not been possible to fully characterise the RNA species making up the other chromatographic peaks due to challenges surrounding signal-to-noise ratio and the increased spectral complexity arising from larger higher molecular weight RNA species and co-eluting sense and antisense strands from the dsRNA products.

Intact mass analysis of RNA synthesised from expression constructs containing three transcriptional terminators has enabled mechanistic insights into the nature of intrinsic transcriptional termination resulting in different length RNA products when multiple transcriptional terminator sequences are used sequentially. The multiple RNA products observed correspond to alternative sites of transcriptional termination with the multiple-terminator RNA expression constructs.

## Conclusions

dsRNA is emerging as a novel sustainable method of plant protection as an alternative to traditional chemical pesticides. The successful commercialisation of dsRNA based biocontrols for effective pest management strategies requires the economical production of large quantities of dsRNA combined with suitable delivery methods to ensure RNAi efficacy against the target pest. In this study, we have optimised the design of plasmid constructs for production of dsRNA biocontrols in *E. coli*. The results show that utilising multiple synthetic transcriptional terminators outside of the dual convergent T7 RNA polymerase promoters in the plasmid construct design results in a significant increase in the amount of dsRNA biocontrol expressed *in vivo* in *E. coli* HT115 cells. Furthermore, the results show that using the same number of *E. coli* cells expressing dsRNA from plasmid constructs containing multiple T7 terminators (3T β*-actin* dsRNA) results in higher RNAi efficacy as measured by insect mortality compared to *E. coli* cells expressing dsRNA from plasmid constructs containing no T7 terminators (NT β*-actin* dsRNA). It is proposed that the increased yield of dsRNA generated from plasmid constructs containing multiple T7 terminators is due to increased termination efficiency *in vivo*, resulting from more efficient utilisation of cellular NTPs and more efficient T7 polymerase activity by preventing RNA synthesis around the DNA plasmid template beyond the sequence of interest. Consistent with this hypothesis is *in vitro* termination efficiency analysis which showed that >90% termination efficiency was achieved using multiple transcriptional terminators compared to 49% for a single terminator. Finally, we have used mass spectrometry to characterise the termination of T7 RNA polymerase and provide further mechanistic insight into T7 polymerase termination using DNA constructs with multiple transcriptional terminators. Using intact mass analysis the results show the most abundant species produced by the T7 RNA polymerase was found to correspond to transcription terminating at the end of the first T7 S terminator sequence within the construct while the next two most abundant species were found to correspond to transcriptional termination at different positions within the larger *rrnB* T1 terminator. The final species characterised through mass spectrometry was found to be terminated once again at the end of the T7 S, the third and final terminator within the construct. These results demonstrate the versatility of the optimised DNA constructs for production of dsRNA that can be potentially used for the production of a wide range of different dsRNA biocontrols across multiple gene targets and can be implemented into other designs if necessary.

## Data availability

The data underlying this article are available in the article and in its online supplementary material or available on request.

## Supplementary data

Supplementary Data are available.

## Funding and Acknowledgements

SJR is a Biotechnology and Biological Science Research Council White Rose DTP iCASE student in collaboration with Syngenta (BB/T007222/1). GRO is an Engineering and Physical Science Research Council student funded from the Institute for Sustainable Food at the University of Sheffield. MJD acknowledges further support from the Biotechnology and Biological Science Research Council (BB/M012166/1).

