## Supplementary Figures for "Optimising the production of dsRNA biocontrols in microbial systems using multiple transcriptional terminators"

### Slide 1
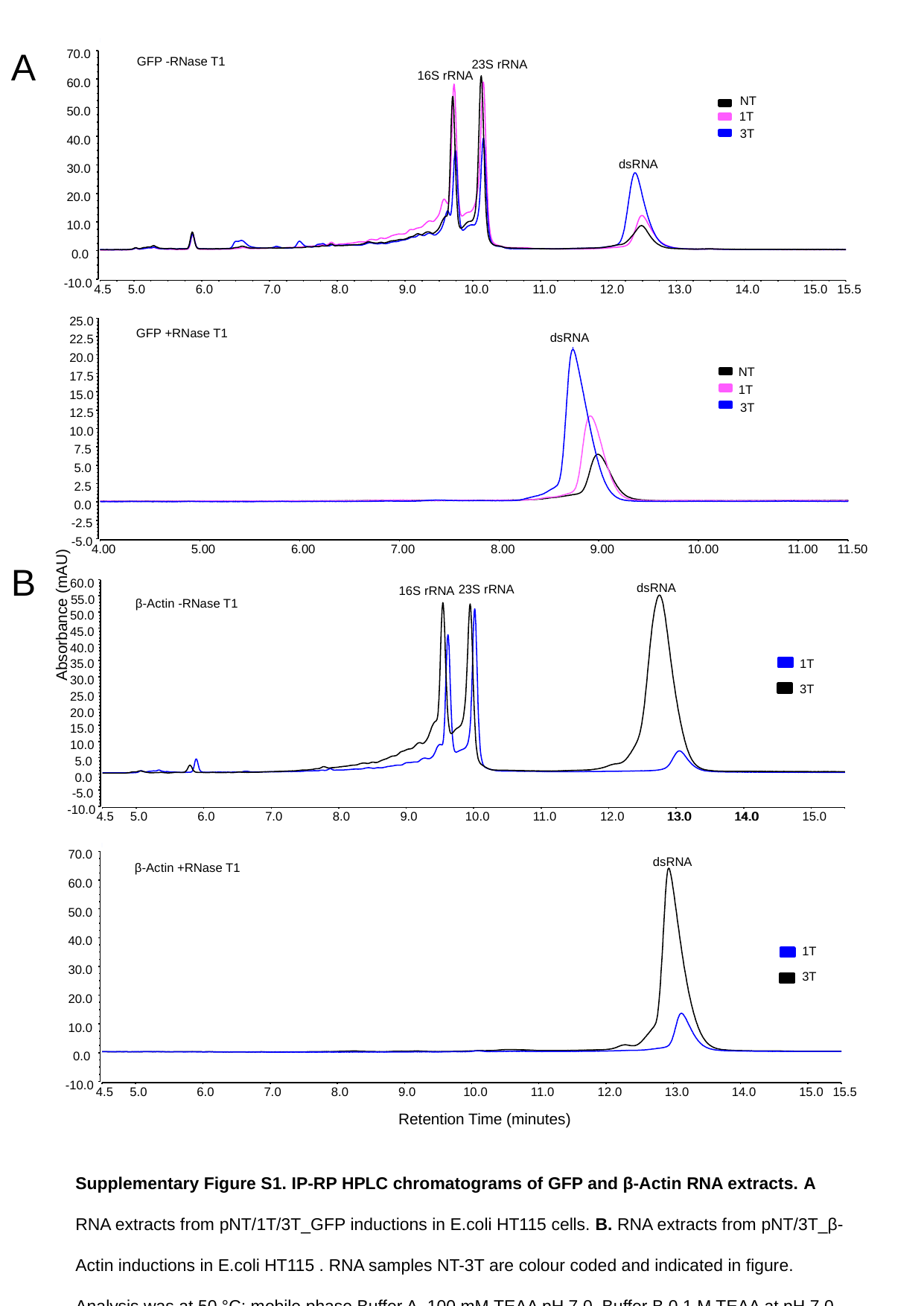

A
70.0
60.0
50.0
40.0
30.0
20.0
10.0
0.0
-10.0
4.5
5.0
6.0
7.0
8.0
9.0
10.0
11.0
12.0
13.0
14.0
15.0
15.5
23S rRNA
16S rRNA
NT
1T
3T
dsRNA
GFP -RNase T1
25.0
22.5
20.0
17.5
15.0
12.5
10.0
7.5
5.0
2.5
0.0
-2.5
-5.0
4.00
5.00
6.00
7.00
8.00
9.00
10.00
11.00
11.50
dsRNA
NT
1T
3T
GFP +RNase T1
Absorbance (mAU)
70.0
60.0
50.0
40.0
30.0
20.0
10.0
0.0
-10.0
4.5
5.0
6.0
7.0
8.0
9.0
10.0
11.0
12.0
13.0
14.0
15.0
15.5
dsRNA
1T
3T
Retention Time (minutes)
dsRNA
60.0
55.0
50.0
45.0
40.0
35.0
30.0
25.0
20.0
15.0
10.0
5.0
0.0
-5.0
-10.0
4.5
5.0
6.0
7.0
8.0
9.0
10.0
11.0
12.0
13.0
14.0
15.0
23S rRNA
16S rRNA
1T
3T
β-Actin -RNase T1
13.0
14.0
β-Actin +RNase T1
B
Supplementary Figure S1. IP-RP HPLC chromatograms of GFP and β-Actin RNA extracts. A RNA extracts from pNT/1T/3T_GFP inductions in E.coli HT115 cells. B. RNA extracts from pNT/3T_β-Actin inductions in E.coli HT115 . RNA samples NT-3T are colour coded and indicated in figure. Analysis was at 50 °C; mobile phase Buffer A, 100 mM TEAA pH 7.0, Buffer B 0.1 M TEAA at pH 7.0 containing 25% acetonitrile, UV absorbance at 260 nm. The gradient use and quantity of RNA analysed can be found in Section XX.

### Slide 2
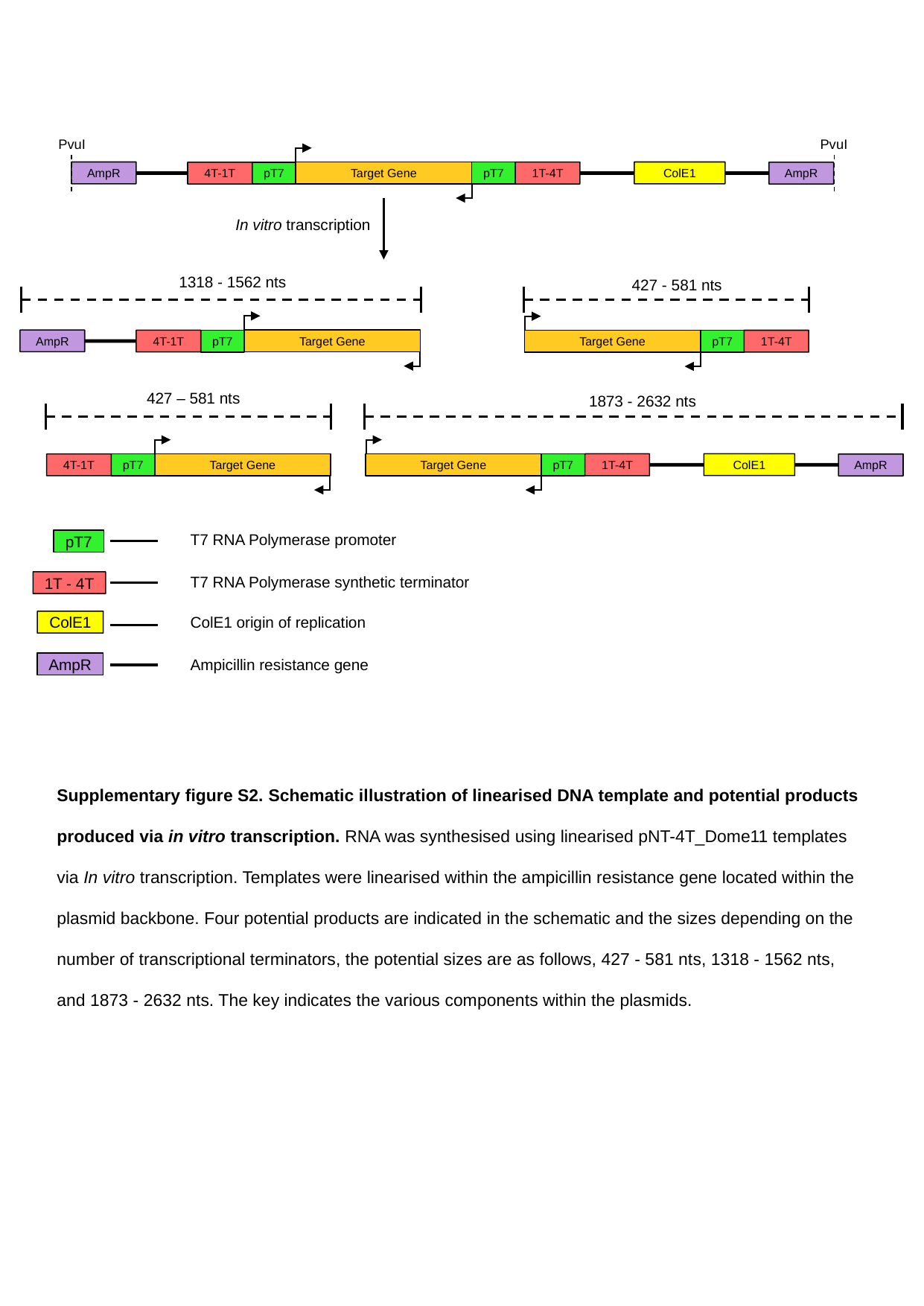

PvuI
PvuI
Target Gene
pT7
pT7
1T-4T
4T-1T
ColE1
AmpR
AmpR
In vitro transcription
1318 - 1562 nts
427 - 581 nts
Target Gene
pT7
4T-1T
AmpR
Target Gene
pT7
1T-4T
427 – 581 nts
1873 - 2632 nts
Target Gene
pT7
1T-4T
ColE1
AmpR
Target Gene
pT7
4T-1T
T7 RNA Polymerase promoter
pT7
T7 RNA Polymerase synthetic terminator
1T - 4T
ColE1 origin of replication
Ampicillin resistance gene
AmpR
ColE1
Supplementary figure S2. Schematic illustration of linearised DNA template and potential products produced via in vitro transcription. RNA was synthesised using linearised pNT-4T_Dome11 templates via In vitro transcription. Templates were linearised within the ampicillin resistance gene located within the plasmid backbone. Four potential products are indicated in the schematic and the sizes depending on the number of transcriptional terminators, the potential sizes are as follows, 427 - 581 nts, 1318 - 1562 nts, and 1873 - 2632 nts. The key indicates the various components within the plasmids.

### Slide 3
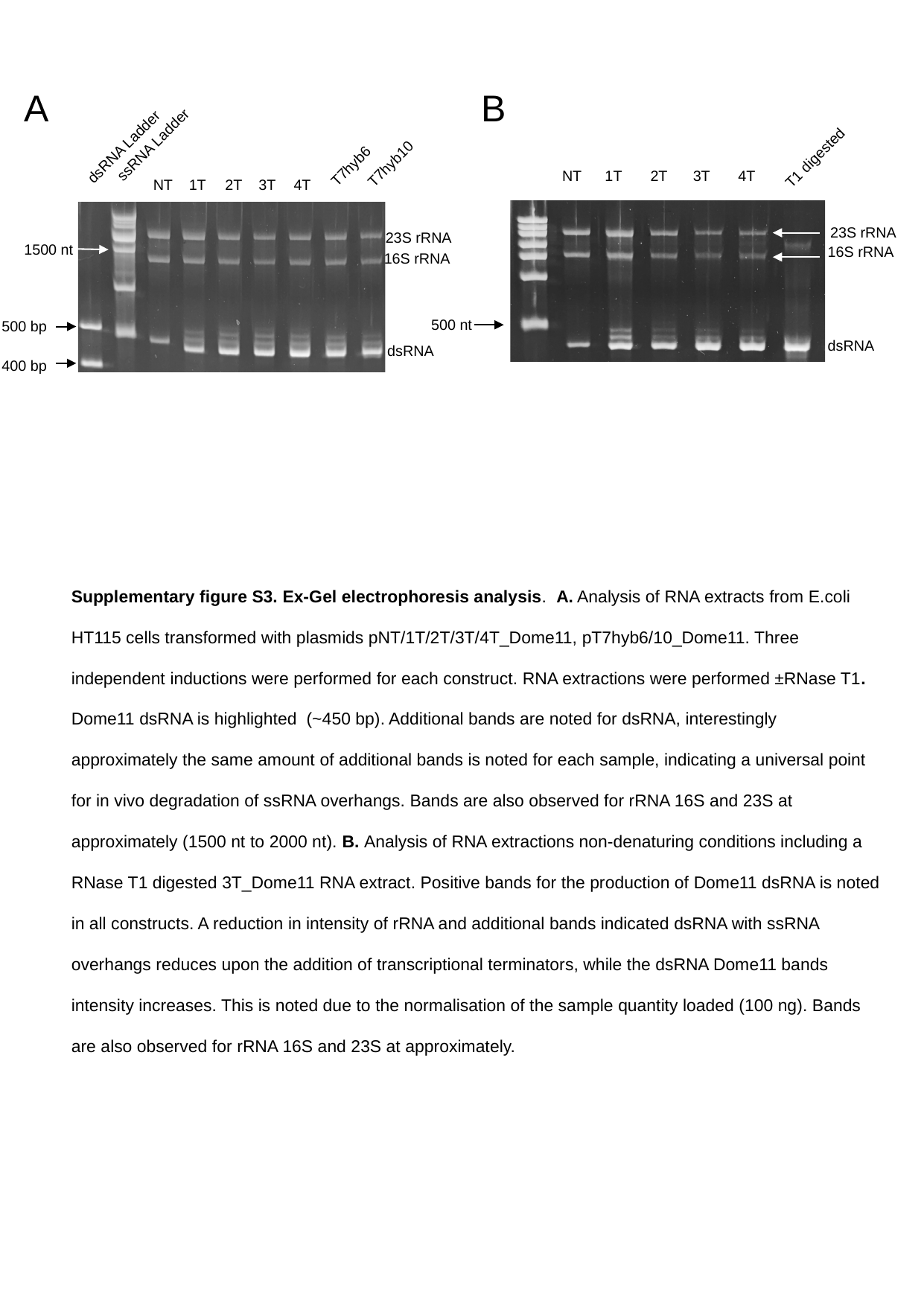

A
B
T1 digested
NT
1T
2T
3T
4T
500 nt
dsRNA
ssRNA Ladder
dsRNA Ladder
T7hyb6
T7hyb10
1T
2T
3T
4T
NT
16S rRNA
500 bp
dsRNA
400 bp
23S rRNA
23S rRNA
1500 nt
16S rRNA
Supplementary figure S3. Ex-Gel electrophoresis analysis. A. Analysis of RNA extracts from E.coli HT115 cells transformed with plasmids pNT/1T/2T/3T/4T_Dome11, pT7hyb6/10_Dome11. Three independent inductions were performed for each construct. RNA extractions were performed ±RNase T1. Dome11 dsRNA is highlighted (~450 bp). Additional bands are noted for dsRNA, interestingly approximately the same amount of additional bands is noted for each sample, indicating a universal point for in vivo degradation of ssRNA overhangs. Bands are also observed for rRNA 16S and 23S at approximately (1500 nt to 2000 nt). B. Analysis of RNA extractions non-denaturing conditions including a RNase T1 digested 3T_Dome11 RNA extract. Positive bands for the production of Dome11 dsRNA is noted in all constructs. A reduction in intensity of rRNA and additional bands indicated dsRNA with ssRNA overhangs reduces upon the addition of transcriptional terminators, while the dsRNA Dome11 bands intensity increases. This is noted due to the normalisation of the sample quantity loaded (100 ng). Bands are also observed for rRNA 16S and 23S at approximately.

### Slide 4
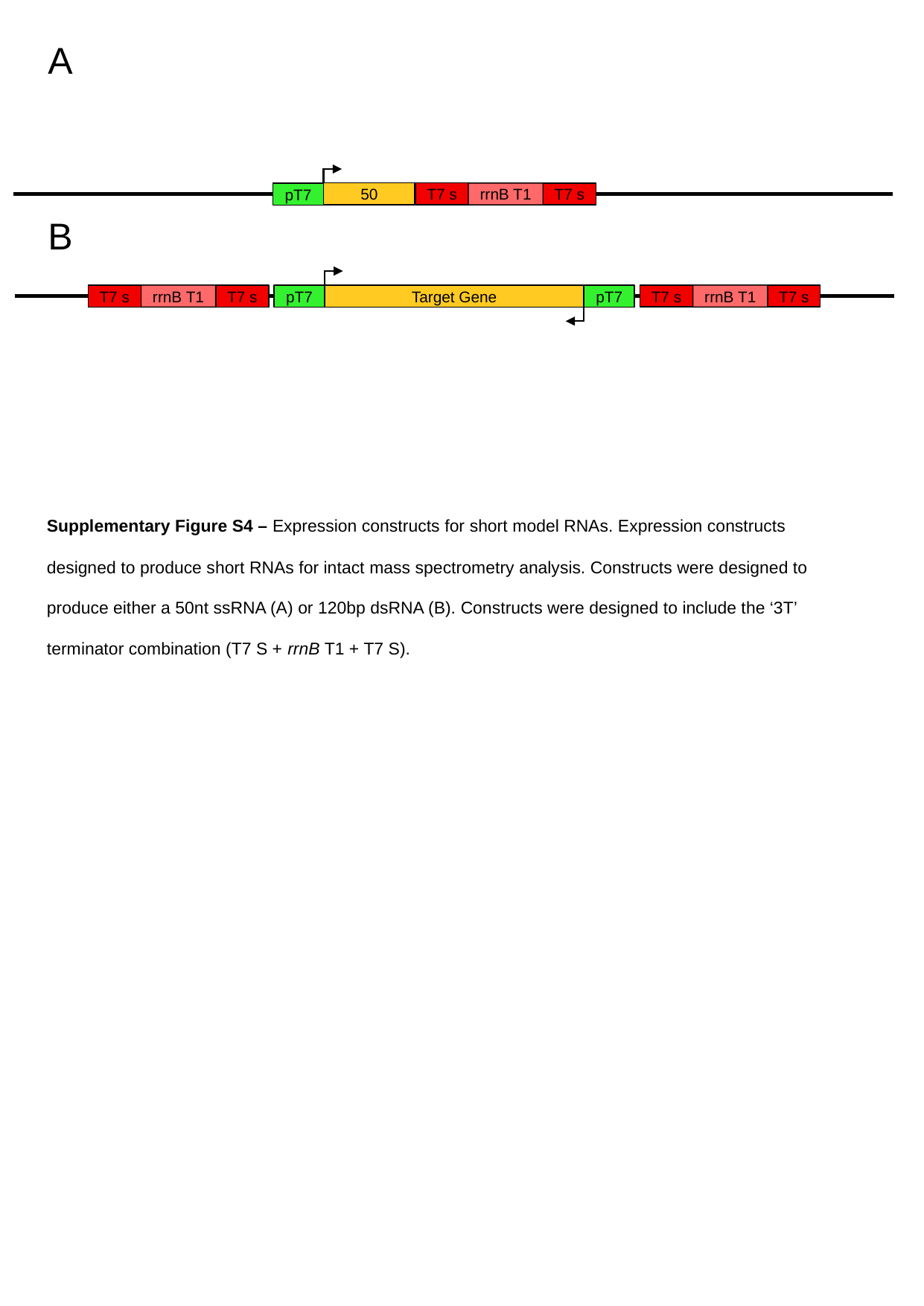

A
T7 s
rrnB T1
50
T7 s
pT7
B
T7 s
rrnB T1
T7 s
rrnB T1
Target Gene
pT7
T7 s
T7 s
pT7
Supplementary Figure S4 – Expression constructs for short model RNAs. Expression constructs designed to produce short RNAs for intact mass spectrometry analysis. Constructs were designed to produce either a 50nt ssRNA (A) or 120bp dsRNA (B). Constructs were designed to include the ‘3T’ terminator combination (T7 S + rrnB T1 + T7 S).

### Slide 5
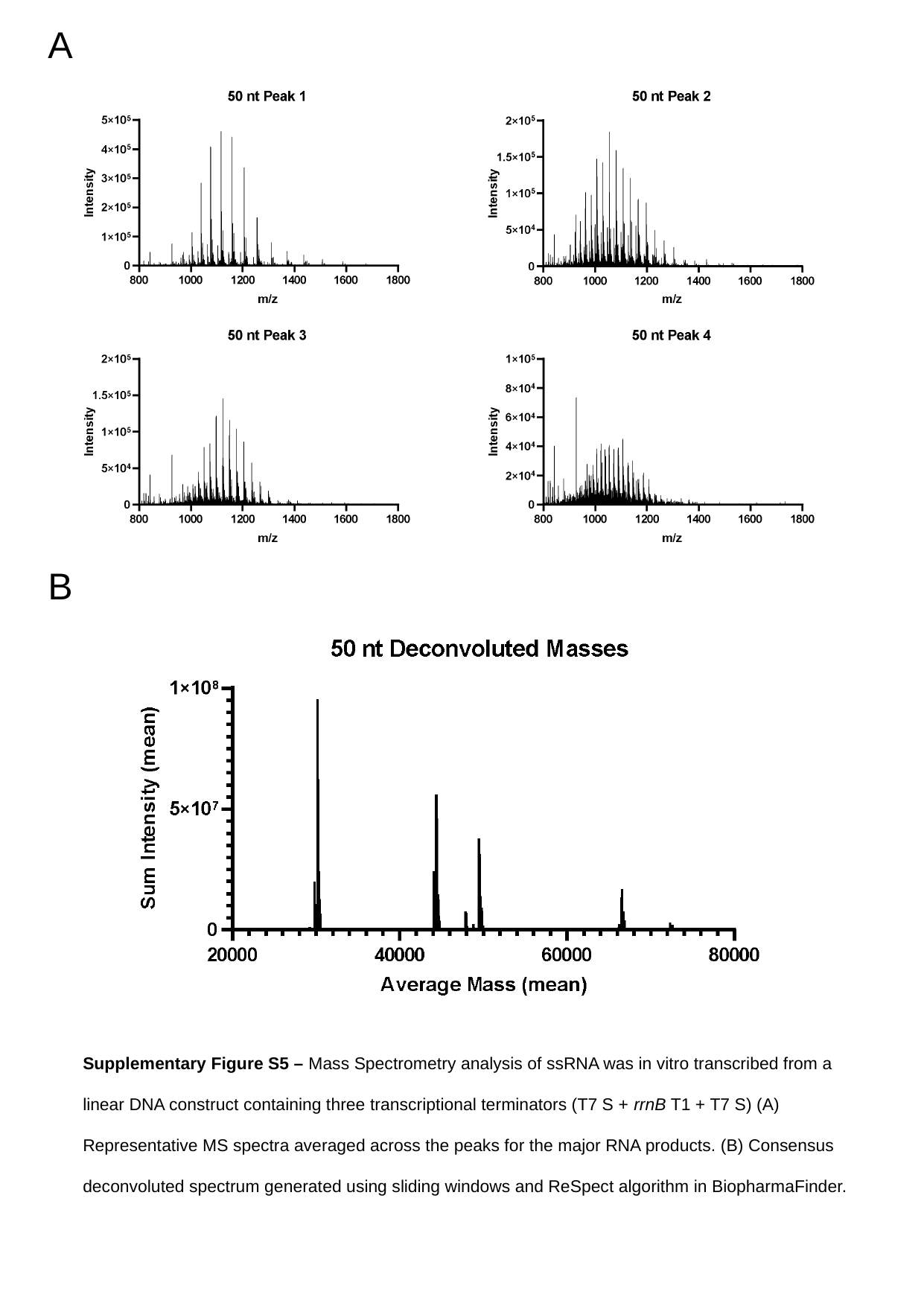

A
B
Supplementary Figure S5 – Mass Spectrometry analysis of ssRNA was in vitro transcribed from a linear DNA construct containing three transcriptional terminators (T7 S + rrnB T1 + T7 S) (A) Representative MS spectra averaged across the peaks for the major RNA products. (B) Consensus deconvoluted spectrum generated using sliding windows and ReSpect algorithm in BiopharmaFinder.

### Slide 6
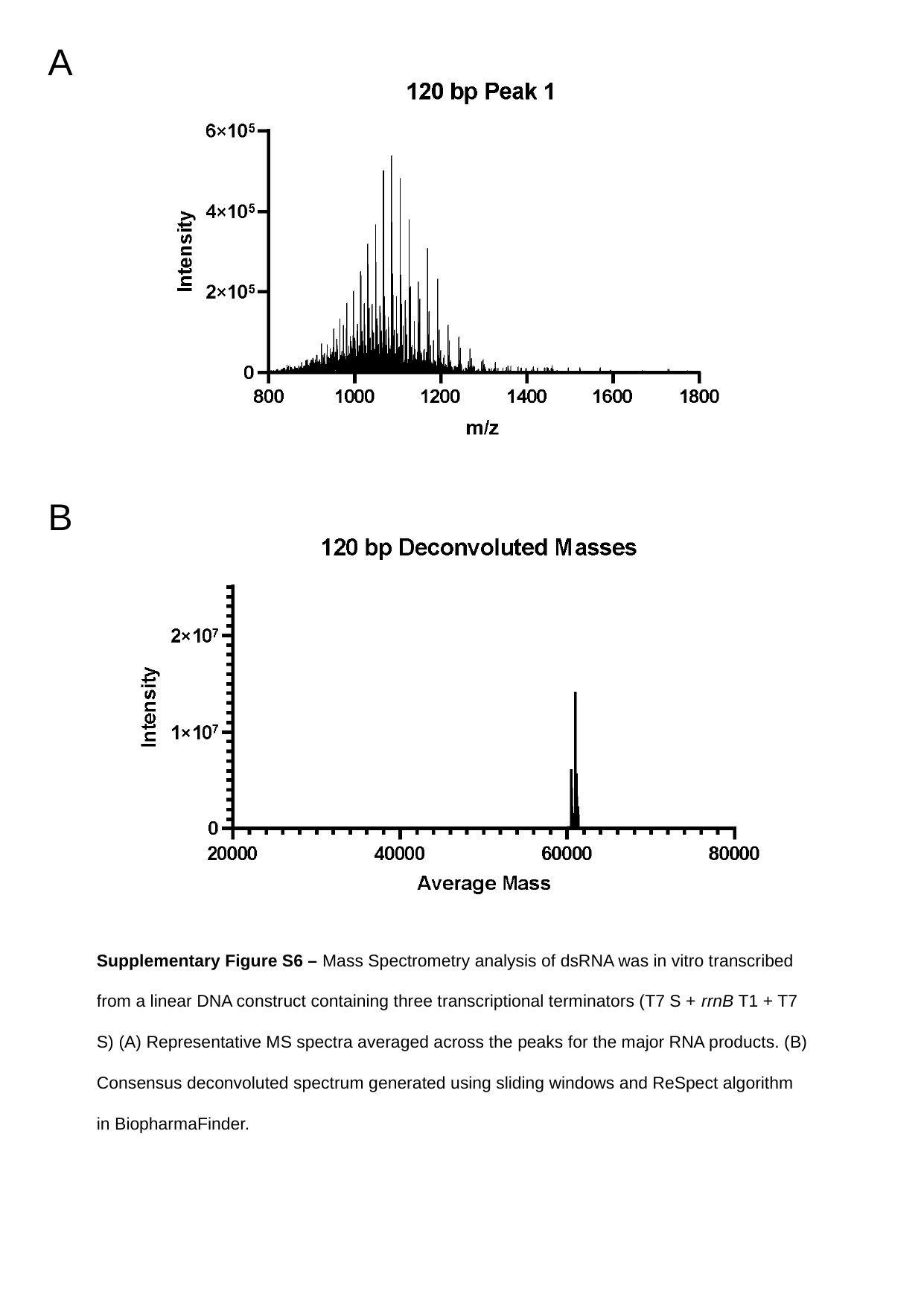

A
B
Supplementary Figure S6 – Mass Spectrometry analysis of dsRNA was in vitro transcribed from a linear DNA construct containing three transcriptional terminators (T7 S + rrnB T1 + T7 S) (A) Representative MS spectra averaged across the peaks for the major RNA products. (B) Consensus deconvoluted spectrum generated using sliding windows and ReSpect algorithm in BiopharmaFinder.

### Slide 7
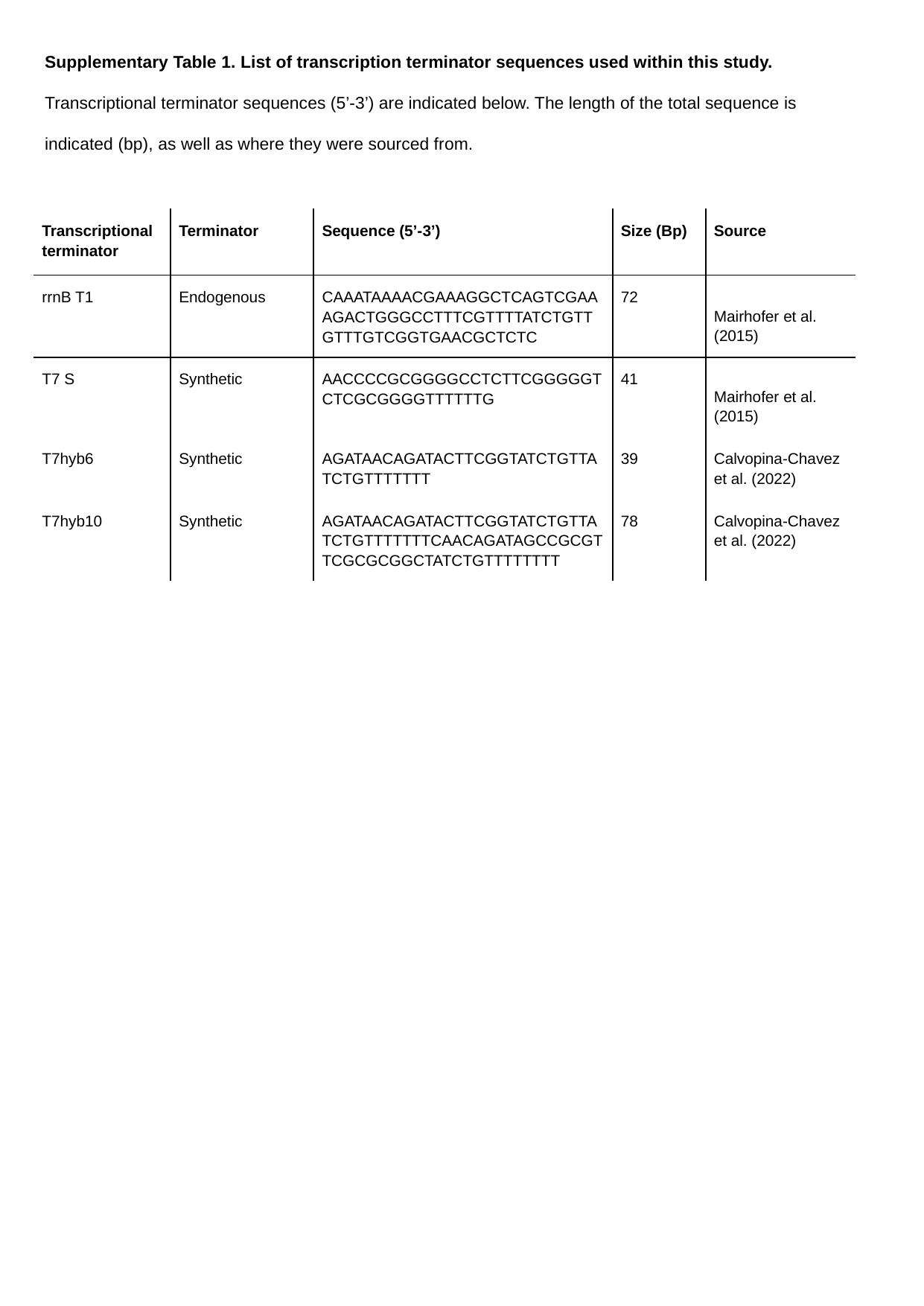

Supplementary Table 1. List of transcription terminator sequences used within this study. Transcriptional terminator sequences (5’-3’) are indicated below. The length of the total sequence is indicated (bp), as well as where they were sourced from.
| Transcriptional terminator | Terminator | Sequence (5’-3’) | Size (Bp) | Source |
| --- | --- | --- | --- | --- |
| rrnB T1 | Endogenous | CAAATAAAACGAAAGGCTCAGTCGAAAGACTGGGCCTTTCGTTTTATCTGTTGTTTGTCGGTGAACGCTCTC | 72 | Mairhofer et al. (2015) |
| T7 S | Synthetic | AACCCCGCGGGGCCTCTTCGGGGGTCTCGCGGGGTTTTTTG | 41 | Mairhofer et al. (2015) |
| T7hyb6 | Synthetic | AGATAACAGATACTTCGGTATCTGTTATCTGTTTTTTT | 39 | Calvopina-Chavez et al. (2022) |
| T7hyb10 | Synthetic | AGATAACAGATACTTCGGTATCTGTTATCTGTTTTTTTCAACAGATAGCCGCGTTCGCGCGGCTATCTGTTTTTTTT | 78 | Calvopina-Chavez et al. (2022) |
